# Reduced biomechanical models for precision-cut lung-slice stretching experiments

**DOI:** 10.1101/2020.07.09.195735

**Authors:** Hannah J. Pybus, Lowell T. Edgar, Reuben D. O’Dea, Bindi S. Brook

## Abstract

Precision-cut lung-slices (PCLS), in which viable airways embedded within lung parenchyma are stretched or induced to contract, are a widely used *ex vivo* assay to investigate bronchoconstriction and, more recently, mechanical activation of pro-remodelling cytokines in asthmatic airways. We develop a nonlinear fibre-reinforced biomechanical model accounting for smooth muscle contraction and extracellular matrix strain-stiffening. Through numerical simulation, we describe the stresses and contractile responses of an airway within a PCLS of finite thickness, exposing the importance of smooth muscle contraction on the local stress state within the airway. We then consider two simplifying limits of the model (a membrane representation and an asymptotic reduction in the thin-PCLS-limit), that permit analytical progress. Comparison against numerical solution of the full problem shows that the asymptotic reduction successfully captures the key elements of the full model behaviour. The more tractable reduced model that we develop is suitable to be employed in investigations to elucidate the time-dependent feedback mechanisms linking airway mechanics and cytokine activation in asthma.

## 1 Introduction

Around 334 million individuals worldwide suffer from asthma and it is estimated that over 250,000 of these people die prematurely each year as a result (Forum of International Respiratory Societies, 2017). Asthma is the most prominent chronic disease amongst youths, affecting over 14% of children globally (Pearce et al., 2007), yet despite its rising prevalence, the cause and onset of asthma remains unknown. Understanding asthma is of vital importance.

Asthma is characterised by inflammation, airway hyperresponsiveness and remodelling (Brightling et al., 2012; Berair et al., 2013). Airway hyperresponsiveness refers to excessive bronchoconstriction (narrowing of the airway) due to rapid contraction of airway smooth muscle (ASM) in response to a relatively low dose of contractile agonist (King et al., 1999; West et al., 2013). Chronic inflammation causes swelling of the airway tissue, narrowing the airway (León, 2017), and resulting in overall restricted pulmonary function. The persistent structural changes due to inflammatory injury repair, airway thickening and scarring constitute airway remodelling (Bossé et al., 2008; Al Alawi et al., 2014). Until recently, airway remodelling has been predominantly attributed to chronic inflammation (Saglani and Lloyd, 2015). Current experimental evidence, however, suggests that bronchoconstriction induced airway narrowing may play a key role in promoting remodelling (Grainge et al., 2011) via activation of the regulatory cytokine, transforming growth factor *β* (TGF-*β*) (Buscemi et al., 2011; Tatler et al., 2011; Wipff et al., 2007). In addition, TGF-*β* has been shown to act as a contractile agonist (Ojiaku et al., 2018). Precision-cut lung-slice (PCLS) stretching experiments are a widely-used ex vivo assay (see, *e.g*., Sanderson (2011) and Tan and Sanderson (2014)) for studying agonist-driven bronchoconstriction and more recently how this links to mechanical activation of TGF-*β*. In particular, Tatler (2016) showed that stretching of PCLS increases TGF-*β* activation. However, results from such studies are difficult to interpret without knowledge of the underlying tissue deformation and stress state.

Airway mechanical behaviour is dominated by ASM and collagen-rich extracellular matrix (ECM). ASM is arranged in fibrous bundles that are oriented helically within the airway wall; this arrangement is thought to enhance the bronchocontractile ability of the smaller airways (Amrani and Panettieri, 2003; Ijpma et al., 2017). The ECM is a delicate mesh of deposited connective tissue and fibrous proteins (Hinz, 2015; Cheng et al., 2016), surrounding ASM cell bundles. In the undeformed state, collagen exhibits a ‘crimped’ structure (Kadler et al., 1996); under strain, these structures straighten to bear load. Varition in their natural lengths mean that ECM is ‘recruited’ successively as strain is increased, imparting a strain-stiffening behaviour to the airway (Wells, 2013).

Early mathematical models of the intact airways, accounting for tension generated by ASM contraction and mechanical properties of the airway wall (*e.g*. Latourelle et al. (2002), Affonce and Lutchen (2006), and Ma and Lutchen (2006)), were based on empirical stress-strain relationships for the whole airway (Lambert et al., 1982, 1993, 1994; Lambert and Wilson, 2005) and the Laplace thin-airway wall approximation (Anafi and Wilson, 2001). With these models, it is not possible to determine tissue stresses within the airway wall nor to separate ASM and ECM contributions to the mechanics. Similarly, early models of airways embedded in parenchyma mimicking PCLS experiments assumed the thin-wall Laplace approximation (Bates and Lauzon, 2007; Khan et al., 2010). Brook et al. (2010) extended these early models to account for multiple airway constituents assuming a finite airway wall thickness for both the intact airway and PCLS models under a plane strain and plane stress approximation respectively. While these allowed for tissue stresses to be determined within the airway wall, the linear elastic framework used meant that predictions were only qualitatively useful. Breen et al. (2012) considered a finite element finite-elasticity model of the PCLS but neglected airway wall thickness and was only concerned with stresses in the lung parenchyma. Following arterial and cardiovascular mechanics (Gasser et al., 2006; Ateshian, 2007; Holzapfel and Ogden, 2010; Hill et al., 2012), a nonlinear elastic single-phase fibre-reinforced airway model assuming finite airway wall thickness under a plane strain approximation was developed by Hiorns et al. (2014). Examples of models that account for mucosal growth, buckling and folding are: Wiggs et al. (1997); Moulton and Goriely (2011); Li et al. (2011); Eskandari et al. (2015). More generally, the mechanics of growth in thin biological membranes is described by Kroon and Holzapfel (2008) and Rausch and Kuhl (2014). Approximate solutions for axisymmetric stretching of thin elastic membranes with traction-free surfaces have previously been determined for isotropic materials (Wong and Shield, 1969; Yang, 1967) but do not account for anisotropy or active contraction. Others consider finite deformations of incompressible rubber membranes with a central solid inclusion and under uniform pressure (Jianbing et al., 2015). Finally, models of agonist driven feedback that are focused on growth and remodelling are presented by Chernyavsky et al. (2014), Aparício et al. (2016), and Hill et al. (2018). However, none of these previous models are suitable descriptions for finite deformation of a thin slice in which the tissue stresses may be determined.

In this study we develop a nonlinear fibre-reinforced biomechanical model of PCLS stretching experiments accounting for ASM contraction in response to agonist exposure and ECM strain-stiffening. Through numerical simulation, we quantify the mechanical stress experienced by the airway wall constituents in response to cyclic stretching that consequently activates TGF-*β*. Additionally, we assess the applicability of two simplifying limits of the model; namely, a one-dimensional membrane representation of the PCLS and an asymptotic reduction in the thin-slice limit (that nevertheless retains a description of axial deformation).

## 2 A biomechanical model of PCLS

The PCLS is a well-established experimental preparation for studying airway reactivity, and corresponding biomechanical response (see, *e.g*., Wang et al. (2008), and Tan and Sanderson (2014)). The key advantage of the PCLS is that vital functional interactions between airways, arterioles, and veins are preserved within the alveolar parenchyma (Sanderson, 2011). PCLS are obtained by inflating human lung tissue with liquid agarose, which is allowed to set and solidify before finely slicing. Stretching of the PCLS is effected by adhering it to a deformable membrane, to which a stretch is applied (Figure 1). In the experiments of Tatler (2016), that form our primary motivation, stretch is applied cyclically, in the form of a sine wave with a 15% amplitude and 0.3Hz frequency, for 24 hours (Tatler, 2016). Stretch is applied with a 5%, 10% and 20% amplitude thereafter.

**Fig. 1.**
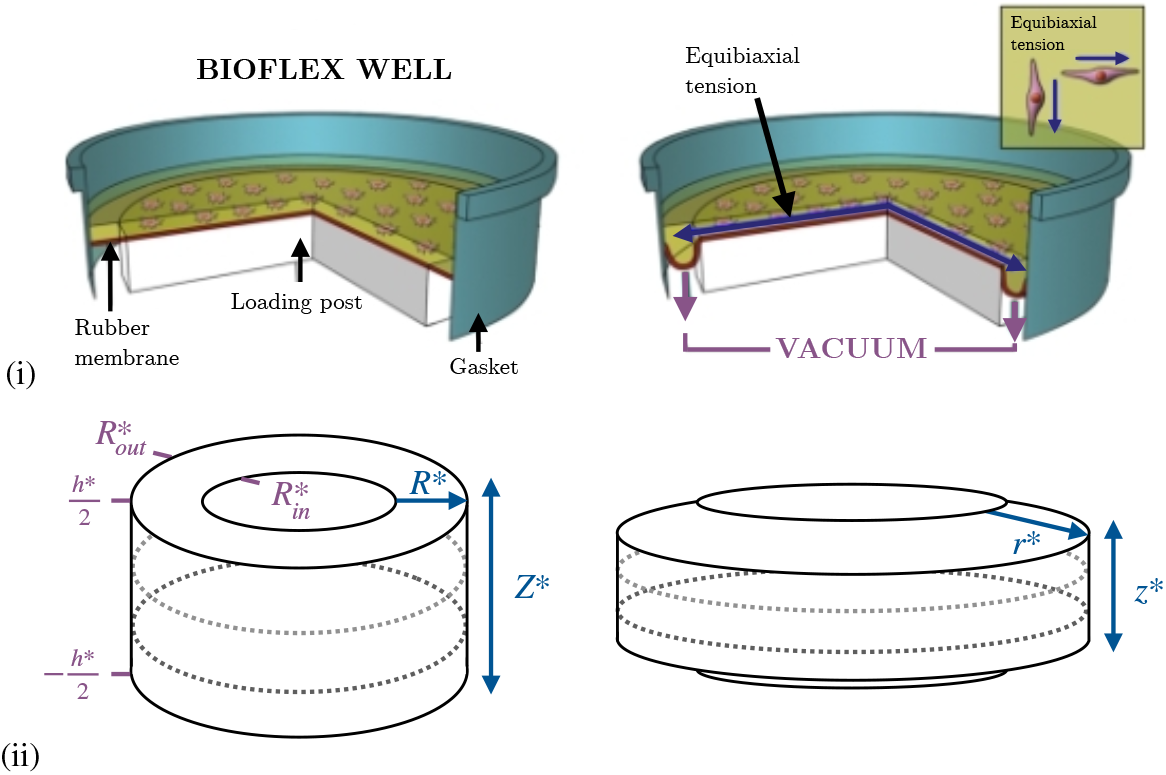
(i) Axisymmetric cyclic stretching of PCLS via the BioFlex method (Tatler, 2016). A PCLS is adhered to a circular deformable rubber membrane and then an axisymmetric cyclic stretch is applied to the membrane (via a vacuum) in order to stretch the attached PCLS. (ii) Dimensional undeformed reference configuration (left) and deformed configuration (right) to illustrate the geometry of an airway modelled within the PCLS. Dotted lines indicate the circumferential fibres representing the active ASM and passive ECM components.

We represent a single airway within the PCLS as a cylinder, whose constituents are modelled via constrained mixture theory (Truesdell and Toupin, 1960; Bowen, 1976; Truesdell and Noll, 1992; Ateshian, 2017). The formulation for obtaining the constitutive mechanical relation for this type of material is given in detail by Humphrey and Rajagopal (2002), Ateshian (2007), and Ateshian and Ricken (2010). Specifically, we consider a saturated multiphase mixture of an active contractile ASM component and a passive ECM component, each modelled as a nonlinear, incompressible, fibre-reinforced hyperelastic material (Holzapfel, 2000), with associated volume fractions *Φ_c_* and *Φ_m_*, respectively. ECM strain-stiffening occurs in the direction of the collagen fibre orientation and accounts for the recruitment of collagen fibres (from a crimped to uncrimped configuration) when stretched (Hiorns et al., 2014). Contractile force generation is assumed to occur in the direction of the ASM bundle orientation and occurs in response to an exogenous agonist and/or active TGF-*β* signalling pathways. Since the duration of the PCLS experiment of interest is significantly less than that of ASM growth or proliferation and ECM deposition, we assume that *Φ_c_* and *Φ_m_* are constant. In addition, for simplicity we neglect the time-dependent feedback between tissue strain and TGF-*β* activation.

### 2.1 Geometry and constitutive assumptions

Following the traditional continuum mechanics approach (Truesdell and Toupin, 1960; Truesdell and Noll, 1992; Holzapfel, 2000) we assume that a common unstressed and unstrained reference configuration applies to each constituent in the airway, in which Lagrangian cylindrical coordinates (*R*\*,Θ,*Z*\*) describe the airway geometry:

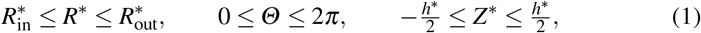

and wherein asterisks denote dimensional quantities, 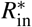 and 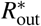 denotes the inner and outer radius and 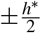 denotes the upper and lower surfaces of the undeformed airway, respectively (see Figure 1 (ii)).

Imposed stretching (herein we use stretch to imply elongation) or contraction of the ASM causes a deformation described by the *deformed configuration* (*r*\*, *θ*, *z*\*). For simplicity, we consider an axisymmetric radial airway stretch, and further assume there is no torsion, so that the deformation is described by

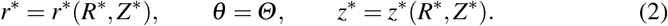

The constituents within the tissue are constrained and therefore also deform axisy-metrically according to (2).

The deformation gradient tensor for the tissue and each constrained constituent within is defined by **F** ≡ **∇x** in the reference configuration and in cylindrical polars is given by

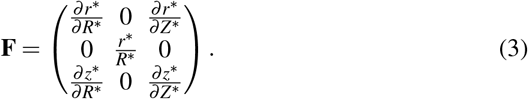

The left and right Cauchy-Green deformation tensors are defined by **B** = **FF**^*T*^ and **C** = **F**^*T*^**F**, respectively. Incompressibility of the tissue is enforced by demanding (Ogden, 2003)

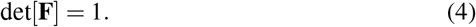

Mechanical anisotropy is imparted to the airway via strain-stiffening of collagen fibres and contractile force generation of ASM bundles (Ogden, 2003), as summarised above. To describe this, we assume that the ASM and ECM constituents within the tissue are associated with a set of fibres orientated circumferentially (Amrani and Panettieri, 2003; Ijpma et al., 2017) with undeformed direction denoted **G**. In the deformed configuration the fibres have direction **g** = **FG**.

Under the above assumptions, the constitutive mechanical law for the airway wall is obtained following the additive approach of Ambrosi and Pezzuto (2012) by introducing an active component, 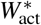, to the passive isotropic, 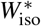, and anisotropic, 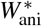, components of the strain-energy function, *W*\*. For each constituent within the tissue, we follow Holzapfel et al. (2000) and define the strain-energy for the ASM and ECM within the tissue as

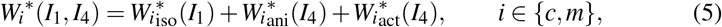

wherein (here and throughout) the subscripts *i* ∈ {*c, m*} denote variables associated with each phase, and the strain energy function for the whole tissue, *W*\* is defined

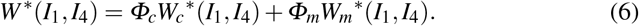

In (5) and (6), *I*_1_ and *I*_4_ denote the first and fourth principle invariants of the left Cauchy-Green deformation tensor, **B**, and are defined by

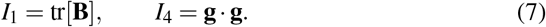

We assume that the isotropic response of the tissue is described via a Neo-Hookean constitutive law, with passive isotropic stiffness 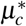 and 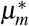 for the ASM and ECM, respectively. It is assumed that collagen fibres within the ECM do not store strain-energy when the airway is not inflated, *i.e*. at low transmural pressures, hence following Holzapfel et al. (2000) we associate an isotropic part of the ECM strain-energy function to the mechanical response of the non-collagenous matrix material. At high transmural pressures the resistance of the tissue to stretch is almost entirely due to anisotropic collagen fibre recruitment within the ECM. To account for strainstiffening, as in Hiorns et al. (2014), we employ the anisotropic model of Holzapfel et al. (2000) with the addition of a Heaviside function so that the collagen fibres are only recruited when stretched. There is no active force contribution from the ECM; however, we include an active component in the ASM strain-energy function. The form of the active component in the Cauchy stress tensor (denoted ***σ***\* and defined below) follows the general form used in Ambrosi and Pezzuto (2012),

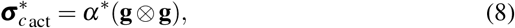

where **g** denotes the direction of the deformed fibres and *α*\* is the active contractile force density (force per unit area) generated by the ASM.

In view of the above, the strain-energy functions for the ASM and ECM components are given by

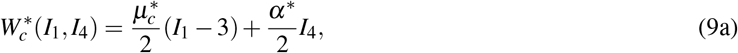

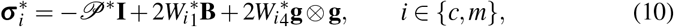

Here, *ω*\* > 0 is a constant parameter defining the passive anisotropic stiffness and accounts for the density of the fibres in the matrix and *ζ* > 0 is a dimensionless constant parameter defining the nonlinear increase in stiffness of the fibres as they deform (Hiorns et al., 2014). The strain-energy function for the whole tissue is given by (6).

The Cauchy stress tensor for each constituent is defined by,

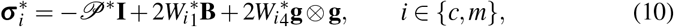

and that for the whole tissue, 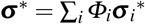. Here, **I** is the identity matrix, the pressure 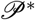 is a Lagrange multiplier included to enforce tissue incompressibility, and 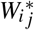 with *i* ∈ {*c, m*} and *j* ∈ {1,4} denote the derivatives of the strain-energy functions with respect to the invariants *I*_1_ and *I*_4_, respectively.

### 2.2 Governing equations and boundary conditions

In mechanical equilibrium, and assuming there are no body forces on the tissue, the balance of linear momentum requires

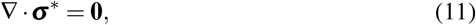

subject to the following boundary conditions.

At the outer radius, we enforce a displacement boundary condition,

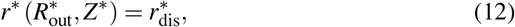

to mimic the axisymmetric stretch imposed on the PCLS via the BioFlex method (Figure 1).

The upper, lower and inner surfaces of the tissue are traction-free such that

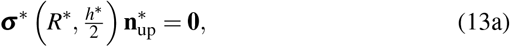

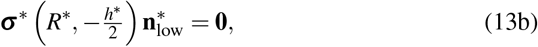

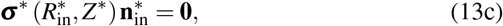

where the unit normals to the upper, 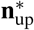, lower, 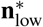, and inner, 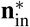, surfaces in the reference configuration are given by:

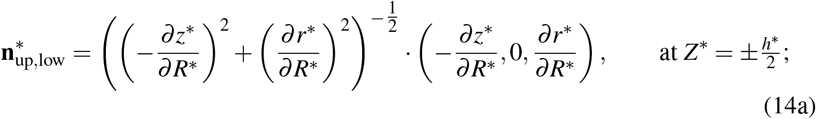

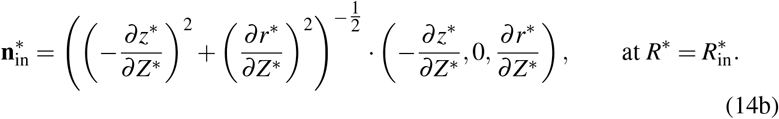

#### 2.2.1 Non-dimensionalisation

We non-dimensionalise the governing equations by introducing the following scalings

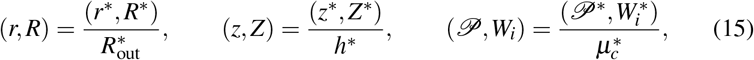

so that the dimensionless undeformed reference configuration is given by

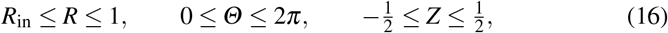

and the deformed configuration is given by

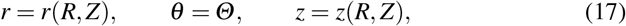

wherein 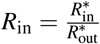 denotes the dimensionless inner radius. Of use in the sequel will be the aspect ratio of the undeformed airway, defined by 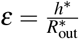.

Under the above definitions, the dimensionless strain-energy functions for each constituent, and the whole tissue are given by

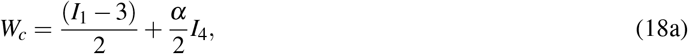

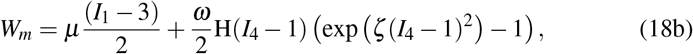

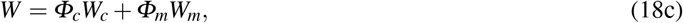

where the dimensionless parameters *μ, ω* and *α* are defined by

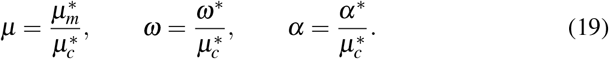

The dimensionless Cauchy stress tenors for each constituent, ***σ***_*i*_, are then obtained from the dimensionless versions of (10); the dimensionless tissue stress is obtained via 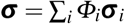.

The dimensionless governing equations (4) and (11) (expressed in terms of the reference configuration) then read

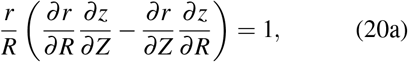

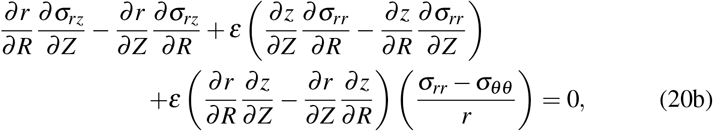

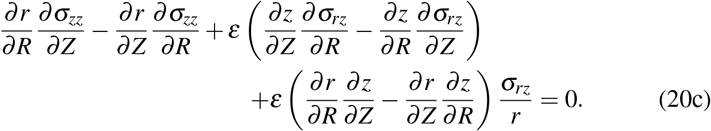

The dimensionless boundary conditions (12) and (13) are given by

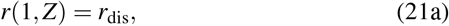

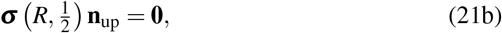

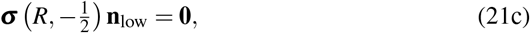

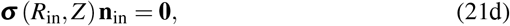

wherein ***n**_up_, **n**_low_* and **n**_*in*_ denote the the dimensionless unit normals to the upper, lower and inner surfaces, respectively.

## 3 Numerical results

Numerical solutions to (20)–(21) are obtained via a finite element method, implemented in the software FEBio (Maas et al., 2012). Figure 2 shows representative results, illustrating the mechanical response of the airway to an imposed radial stretch in the absence (passive case) and presence of active contraction. Details of convergence tests are given in Appendix A and parameter values are given in the relevant figure captions (see also Table 1 in Appendix C).

**Fig. 2.**
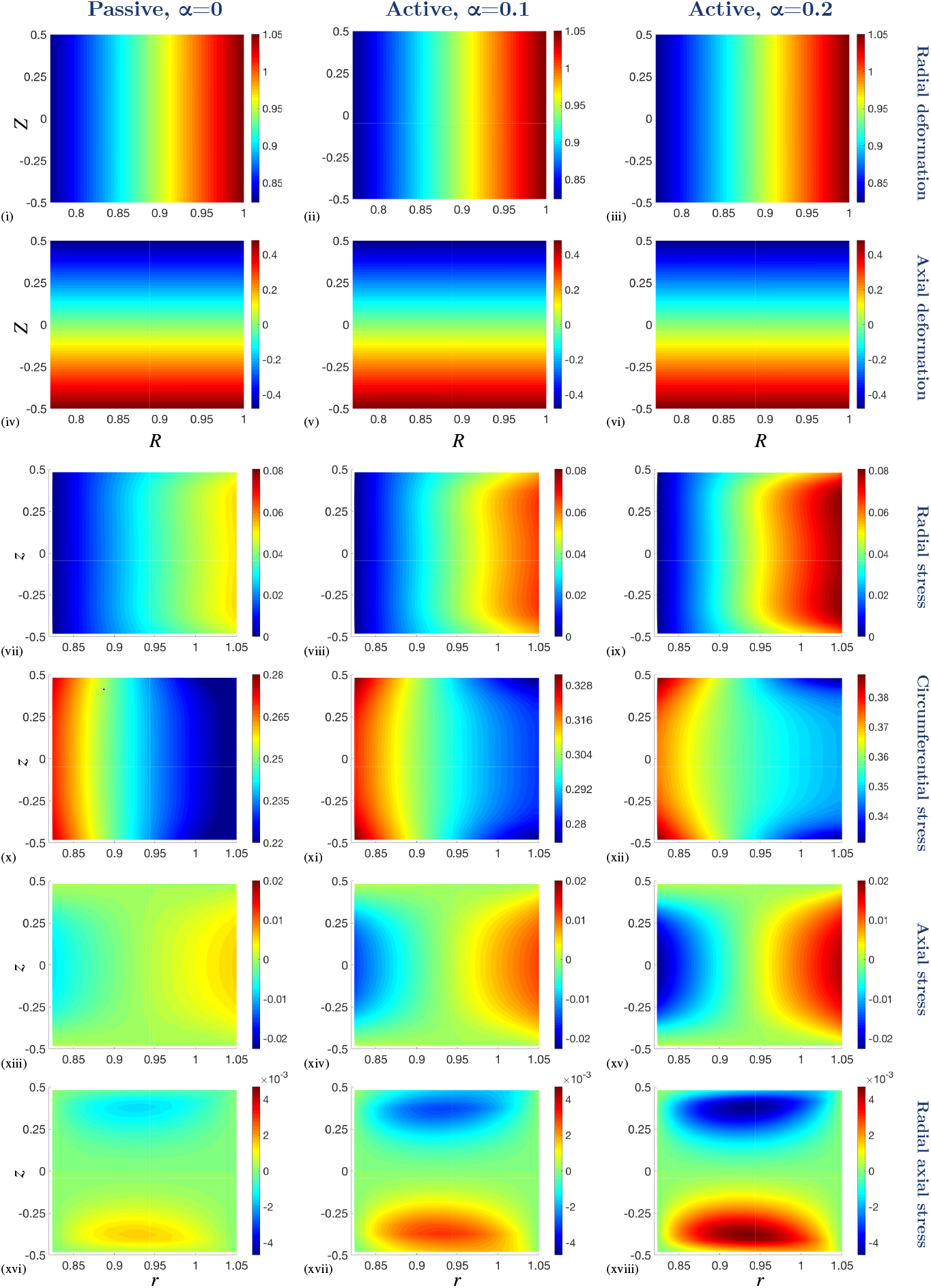
Numerical results from the full thickness model (*ε* = 1) in various states of contraction (*α* = 0, *α* = 0.1 and *α* = 0.2) and with a 5% fixed stretch applied at the outer boundary of the PCLS. (i)–(iii) Radial deformation, *r*, and (iv)–(vi) axial deformation, *z*, plotted over the undeformed configuration, (*R, Z*). Cauchy stress components (vii)–(ix) *σ_rr_*, (x)–(xii) *σ_θθ_*, (xiii)–(xv) *σ_zz_* and (xvi)–(xviii) *σ_rz_* plotted over the deformed configuration, (*r, z*). The parameter values are provided in Table 1 in Appendix C.

**Table 1.**
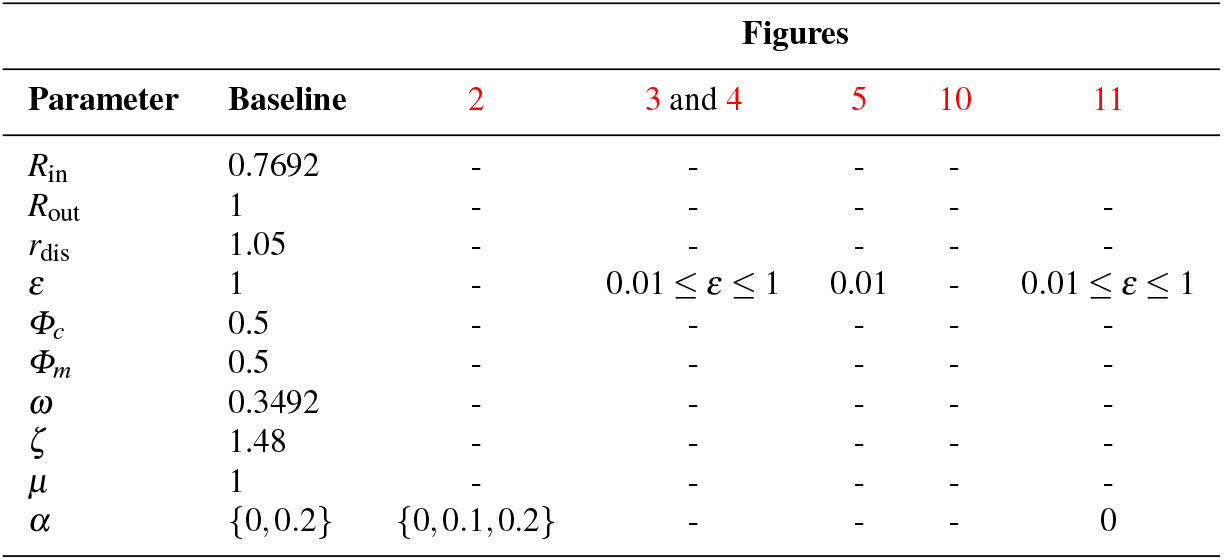
Table of dimensionless parameter values used in Sections 3 and 4.3, and appendices A and B. Dashes denote the given baseline value. Where multiple values are given, the value used for each plot in the figure is specified in the figure’s legend. Where ranges of parameter values are given, the parameter is varied and takes values in the range that are specified in the figure’s legend.

In both the passive and active contraction cases we observe that the radial deformation, *r*, decays linearly with undeformed radius, *R*, but remains uniform axially (Figures 2 (i)–(iii)). As required by the incompressibility of the material, the airway thins as it is stretched (Figures 2 (iv)–(vi)).

The mechanical stress within the tissue displays significant spatial heterogeneity (*e.g*. Figure 2 (xviii)). Moreover, we observe that while the deformation of the airway is qualitatively similar in the passive and active contraction cases (*cf*. Figures 2 (i), (iii)), there are distinct qualitative and quantitative differences in the stress state between these regimes (*cf*. Figures 2 (xvi), (xviii)). In particular, there is an increased and exaggerated heterogeneous stress distribution in the presence of active contraction. Furthermore, the axial dependence of these heterogeneous stress distributions increases with increased active contraction and is highlighted by the circumferential stress distribution, *σ_θθ_* (Figure 2 (xii)).

In each case we observe increased radial stress, *σ_rr_*, at the outer boundary (in the direction of the prescribed stretch), with the stress at the inner wall remaining approximately zero in each case (Figures 2 (vii)–(ix)). Similarly, tissue contractility significantly influences the circumferential stress, *σ_θθ_*, as is to be expected, since the generated contractile stress acts in the direction of the circumferential fibres embedded in the airway (Figures 2 (x)–(xii)). Moreover, we see that the circumferential stress is higher at the inner radius than at the outer radius (Figures 2 (x)–(xii)).

The thinning and stretching of the PCLS under the imposed stretch is reflected in the distributions of the axial, *σ_zz_*, and shear, *σ_rz_*, stresses, with an order of magnitude increase observed in the axial stress in the presence of contraction (Figures 2 (xiii)–(xviii)). The positive (tensile) axial stress at the outer radius and negative (compressive) axial stress at the inner radius reflects the relative thickening at the inner radius compared with that at the outer. The shear stress is positive at the lower surface and negative at the upper surface reflecting the relatively increased displacement of material radially and downward at the upper (and upward at the lower) surface.

The preceding results correspond to an airway of thickness comparable to its outer radius (in particular, we set *ε* = 1). The typical thickness for a PCLS is in the range of 100-250μm and a typical airway radius is in the range of 1-5mm, giving 0.02 < *ε* < 0.25. Motivated by this, we investigate the dependence of the model behaviour on the PCLS thickness by varying the aspect ratio *ε*. We consider the passive, *α* = 0, and active contraction case, *α* = 0.2, in Figures 3 and 4. Here, we reduce *ε* from *ε* = 1 to *ε* = 0.01 in the direction of the black arrows. Note that the variables are plotted as a function of deformed radius at the undeformed axial centre of the PCLS, *i.e*. at *Z* = 0 (Figure 3). This is true in all cases apart from the axial deformation, *z*, which we plot as a function of radius at the undeformed upper surface of the PCLS, *i.e*. at 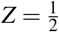, in order to illustrate the thinning of the PCLS in response to stretch (Figures 3 (iii), (iv)). The results illustrating axial dependence are all plotted over the thickness of the PCLS at the undeformed radial midpoint *R* = *R*_mid_ (Figure 4).

**Fig. 3.**
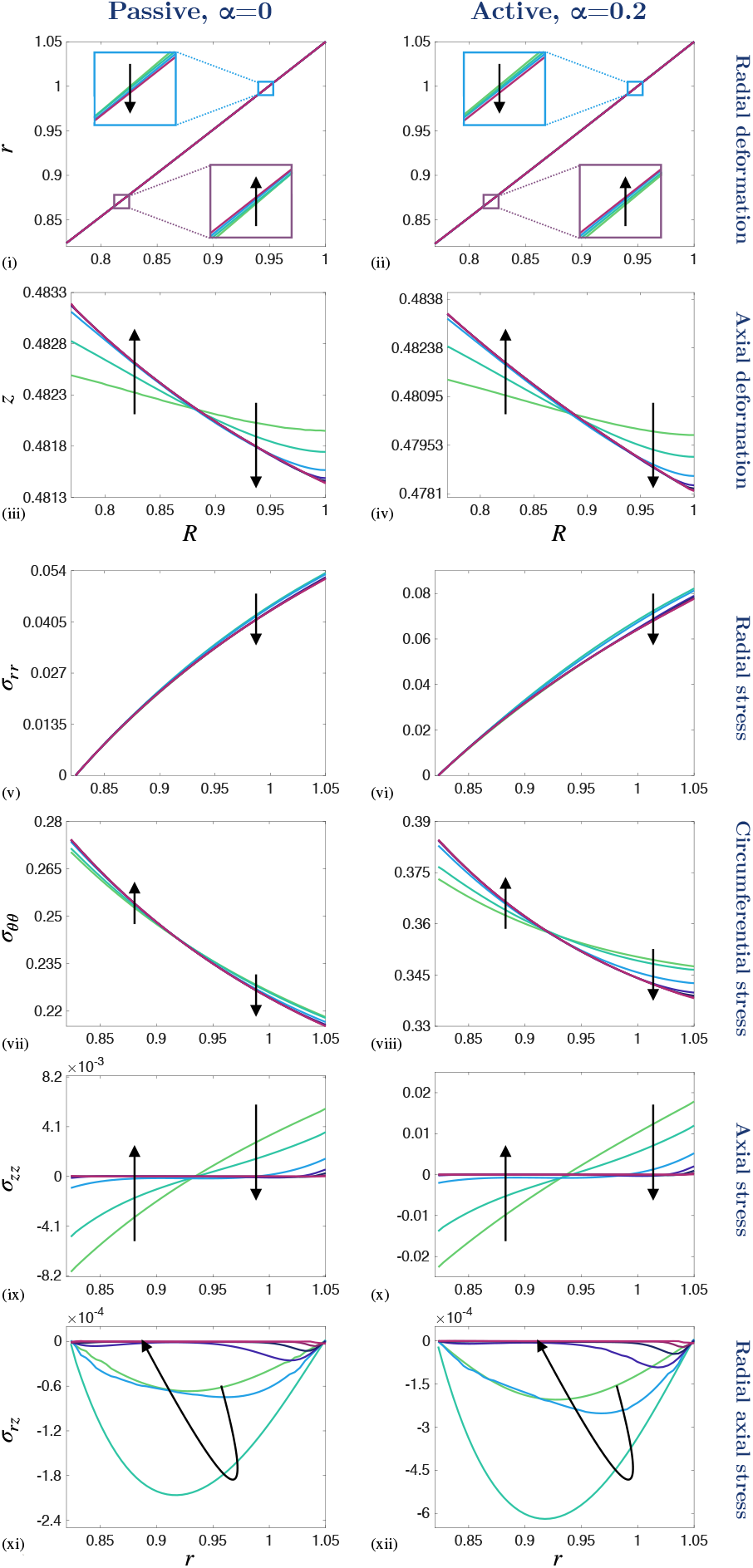
The effect of reducing the PCLS aspect ratio, *ε*, on the deformation and stress distributions across the radius of the airway wall. (i)–(ii) Radial deformation, *r*(*R*, 0), and (iii)–(iv) axial deformation, 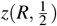, plotted over the undeformed radius, *R*. Cauchy stress components (v)–(vi) *σ_rr_*, (vii)–(viii) *σ_θθ_*, (ix)–(x) *σ_zz_* and (xi)–(xii) *σ_rz_* plotted over the deformed radius, *r*, at *Z* = 0. A 5% fixed stretch is applied in the passive, *α* = 0, and active contraction case, *α* = 0.2. The aspect ratio decreases in direction of black arrows for *ε* ∈ {1,0.5,0.25,0.1,0.05,0.025,0.01} and the remaining parameter values are provided in Table 1 in Appendix C.

**Fig. 4.**
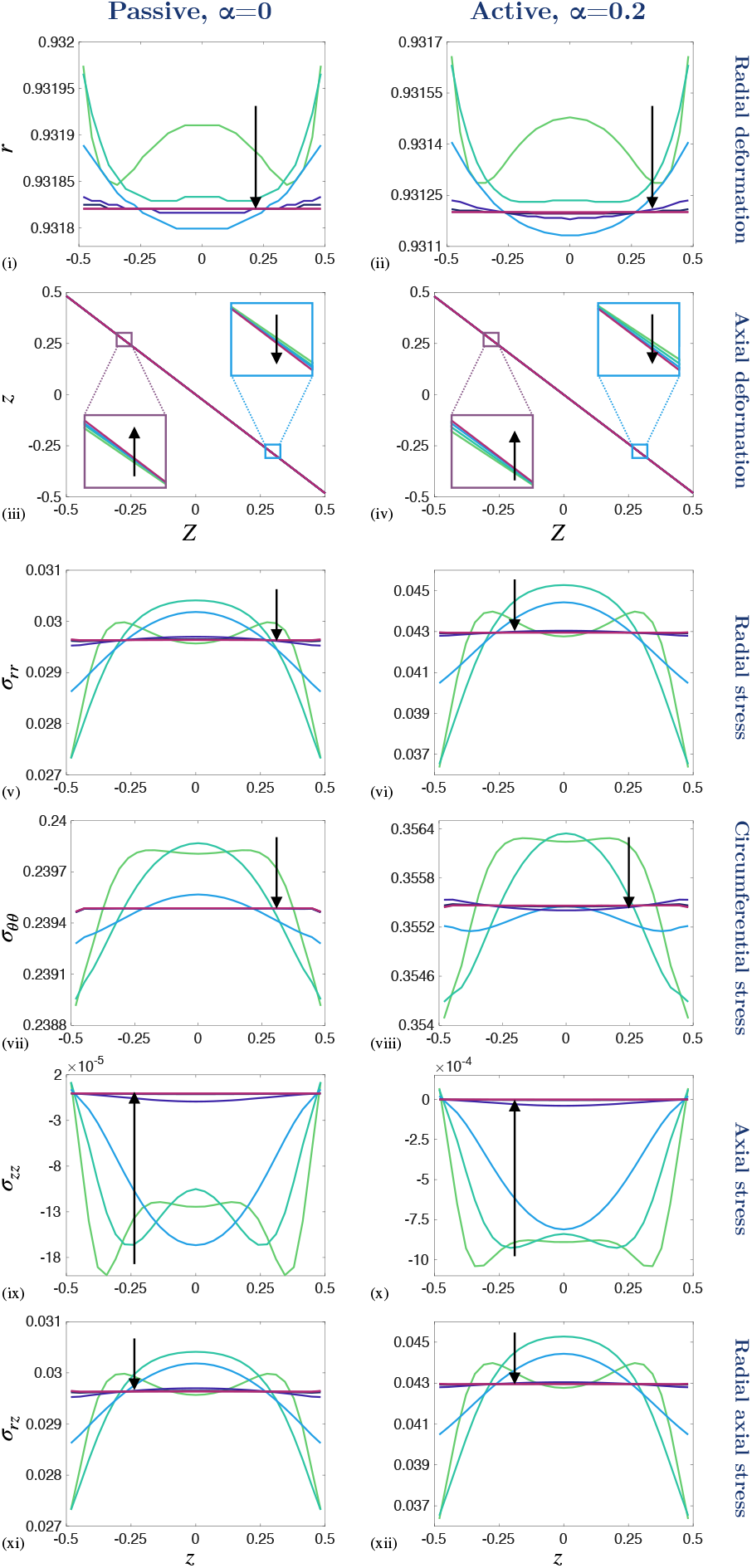
The effect of reducing the PCLS aspect ratio, *ε*, on the deformation and stress distributions through the axial thickness of the PCLS. (i)–(ii) Radial deformation, *r*, and (iii)–(iv) axial deformation, *z*, plotted over the undeformed thickness, *Z*. Cauchy stress components (v)–(vi) *σ_rr_*, (vii)–(viii) *σ_θθ_*, (ix)–(x) *σ_zz_* and (xi)–(xii) *σ_rz_* plotted over the deformed thickness, *z*, at *R* = *R*_mid_. A 5% fixed stretch is applied in the passive, *α* = 0, and active contraction case, *α* = 0.2. The aspect ratio decreases in direction of black arrows for *ε* ∈ {1,0.5,0.25,0.1,0.05,0.025,0.01} and the remaining parameter values are provided in Table 1 in Appendix C.

In both the passive and active contraction cases, the radial deformation at the axial centre line varies linearly with *R* and remains approximately invariant with *ε* (Figures 3 (i), (ii); insets highlighting the very slight variation with *ε*) and with correspondingly little change in the radial stress (Figures 3 (v), (vi)). Conversely, the nearuniform thinning of the airway, observed in Figures 2 (iii), (iv), becomes marginally more exaggerated as *ε* is reduced and the PCLS thins more at the outer radius than at the inner radius. Similarly, the circumferential stress increases at the inner radius and decreases at the outer radius as *ε* reduces (Figures 3 (vii), (viii)). On the other hand, we observe that the heterogeneous axial and shear stress distributions decay to zero in Figures 3 (ix)–(xii).

The deformation and stress variation through the axial thickness is shown in Figure 4. Here we observe only a weak dependence of the radial deformation and stresses on *Z*, that decays to uniformity with decreasing *ε*. In particular, the axial stress decays to zero with *ε* (Figures 4 (ix), (x)). In contrast, the axial deformation remains approximately linear in *Z* as *ε* decreases (Figures 4 (iii), (iv); insets highlighting the very slight variation with *ε*). In the active contraction case, the above described features persist, but the variations in deformation and associated stresses are exaggerated quantitatively (*cf*. Figures 3, 4).

## 4 Model reduction

Guided by the observations in Section 3, in this section we consider analytical simplifications of the biomechanical model (20)–(21). Firstly, in Section 4.1, we adopt a membrane model, following Wong and Shield (1969), that allows reduction to one spatial dimension. Subsequently in Section 4.2, we consider an asymptotic approach to obtain a reduced model describing the leading order PCLS deformation in the thin-PCLS-limit. In Section 4.3, we address the suitability of each approximation by comparing them to the full biomechanical model simulated in FEBio (Maas et al., 2012).

### 4.1 Membrane model

In this section we simplify the biomechanical model (20)–(21) by assuming that the PCLS behaves as an elastic membrane, in which case we neglect *Z* dependence and set *z* = *Z* so that there is no change in the axial thickness of the PCLS upon deformation. This membrane description has previously been used by Wong and Shield (1969) to approximate an axisymmetric stretch of an isotropic sheet; however, Wong and Shield (1969) find that the approximation breaks down when the sheet has an edge which is traction-free. The determinant of the deformation gradient approaches zero in the close vicinity of the traction-free edge and a singularity appears in the governing equations when the material is incompressible. Similarly, although we consider an anisotropic material with active contractile force generation, we find inconsistencies with the governing equations and the prescribed boundary conditions when the thickness of the PCLS is fixed. Specifically, the traction-free boundary conditions on the upper, lower and inner surfaces of the PCLS cannot be satisfied simultaneously, whilst preserving incompressibility, without the PCLS changing in thickness. Therefore, enforcing torsion-free axisymmetry as previously, we reduce the description of the PCLS to one spatial dimension and omit the traction-free boundary conditions on the upper and lower surfaces. The Cauchy stress, ***σ***, and pressure, 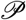, are functions of *R* only, satisfying

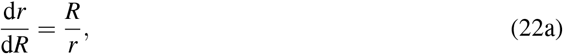

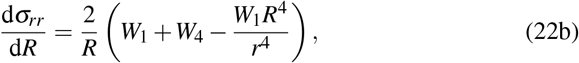

subject to the displacement outer boundary condition (21a),

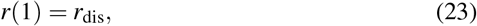

and the free boundary condition at the inner radius (21d) (ommiting (21b)–(21c)),

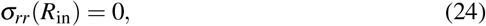

and where the radial and circumferential stresses are constitutively defined as

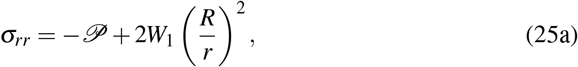

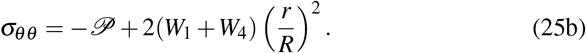

Integrating (22a) with respect to *R*, and imposing (23), we obtain

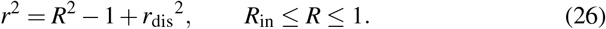

Subsequently, integrating (22b) with respect to *R* and applying the zero radial stress condition at the inner boundary (24) gives

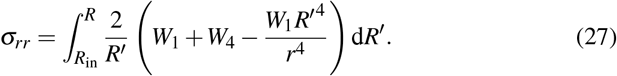

In order to obtain *σ_θθ_* (25b), we require the pressure, 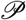; combining (25) and (27) provides

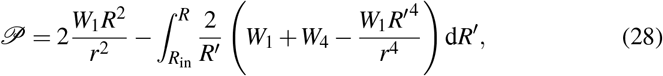

and the constituent and total tissue stress follow directly.

### 4.2 Thin-PCLS-limit

In this section, motivated by the typical geometry of the PCLS (Section 3), we consider the limit 0 < *ε* ≪ 1, so that the thickness of the PCLS is small in comparison to a typical airway radius. Correspondingly, and in view of our numerical results in Section 3 (in particular, Figure 4 where we observe *r* becomes independent of *Z* for 0 < *ε* ≪ 1), we seek expansions of the form

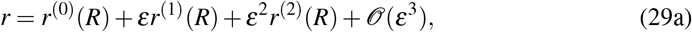

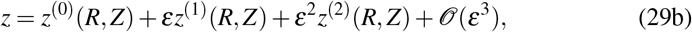

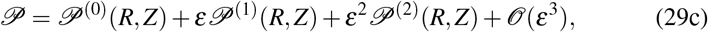

adopting corresponding notation for the strain-energy functions where necessary and assuming that *Φ_c_, Φ_m_, ω, ζ* and *α* all remain 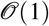 constants. We pause to highlight that the more general expansion, for which *r* = *r*(*R*, *Z*), and the leading term for 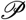 is 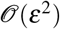 (to obtain the proper leading order balance in the Cauchy stress), can be reduced to that shown in (29) (see Appendix B) and so we adopt this from the outset for brevity.

At leading order, the governing equations (20) read

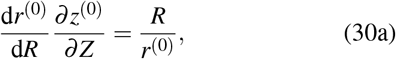

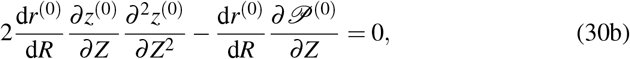

subject to the displacement boundary condition at the outer radius (21a),

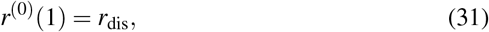

and the following free boundary conditions at the upper, lower and inner surfaces of the PCLS (21b)–(21d):

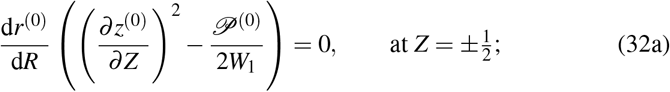

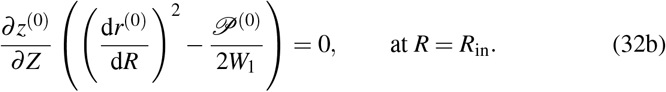

Equation (30a) and the boundary conditions (32a) provide

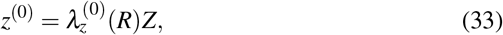

where the arbitrary function of *R* arising from the integration of (30a) vanishes due to axial symmetry in *z*^(0)^ about the axial centre line, *Z* = 0. Furthermore, the equation (32b) requires 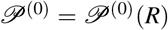. In view of which, together with the boundary conditions (32), we obtain:

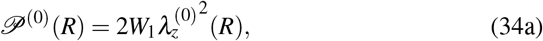

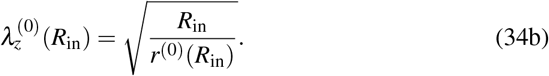

At 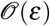 the linear momentum equations (20b)–(20c) read

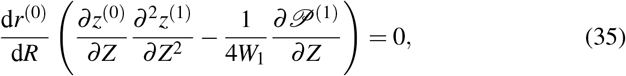

and the boundary conditions (21) provide

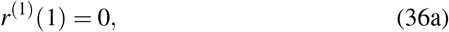

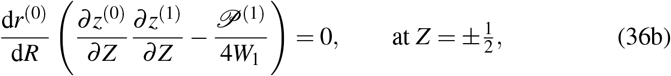

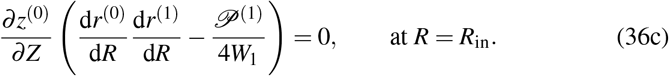

Inspection of equation (36c) shows that 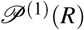 (*R*). In view of which, the boundary conditions (36b) and (36c) provide

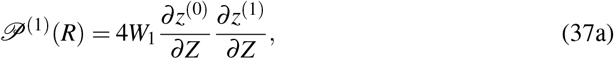

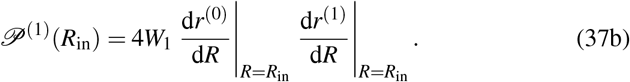

To summarise, we have reduced the problem to two leading order variables; the radial deformation, *r*^(0)^(*R*), and the axial stretch, 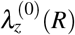, and obtained the governing equation (30a) and the boundary conditions (31) and (34b). However, we require a second governing equation to determine the two variables, *r*^(0)^(*R*) and 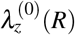. Therefore, we consider the 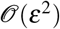 momentum equations; (20b) reads

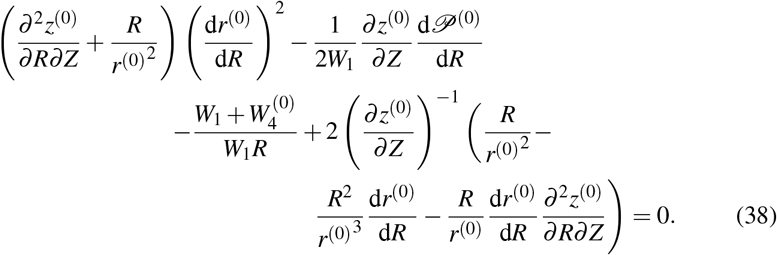

Equation (38) closes the leading order problem. Equation (20c) introduces higher order terms which are not of interest for the leading order problem and is therefore not needed here.

Substituting (33) and (34a) into (38) provides an equation for 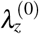 and hence, together with (30a), we obtain the following pair of coupled ODEs:

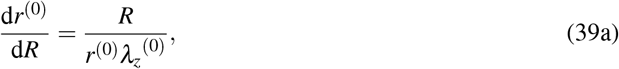

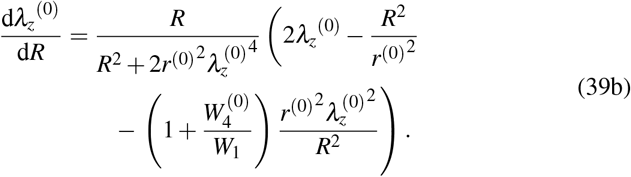

Together with the boundary condition (31) and the relation (34b), this provides a boundary value problem that describes the leading order radial and axial deformation. From this the leading order Cauchy stress components for the whole tissue and each of the constituents follow directly. We note that the boundary condition (34b) on 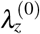 is posed at the (unknown) deformed inner radius. We therefore solve (39) numerically by treating *r*^(0)^(*R*_in_) as a shooting parameter and seek the solution set 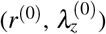 at *R*_in_ that satisfies (31), (34b) and (39).

### 4.3 Suitability of approximations

In this section we compare numerical simulations of the full model (equations (20) with boundary conditions (21)), with those of the membrane model (equations (22) with boundary conditions (23)–(24)), and the thin-PCLS-limit (equations (39) with boundary conditions (31) and (34b)) to demonstrate their validity.

We plot the radial deformation, *r*, axial deformation, *z*, and the corresponding stresses, ***σ***, obtained in all three models, both in the presence and absence of active contractile force in Figure 5. Results from the full and thin-PCLS models in Figure 5 are plotted as functions of radius at the undeformed axial centre line (*Z* = 0), apart from the axial deformation, *z*, which we plot as a function of radius at the undeformed upper surface of the PCLS 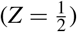, in order to illustrate the thinning of the PCLS in response to stretch. Those from the membrane case, however, do not depend on *Z*; for illustrative purposes, we plot *z* fixed at 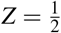 in order to emphasise the thinning of the PCLS (relative to the reference configuration) that is displayed by the full and thin-PCLS models.

**Fig. 5.**
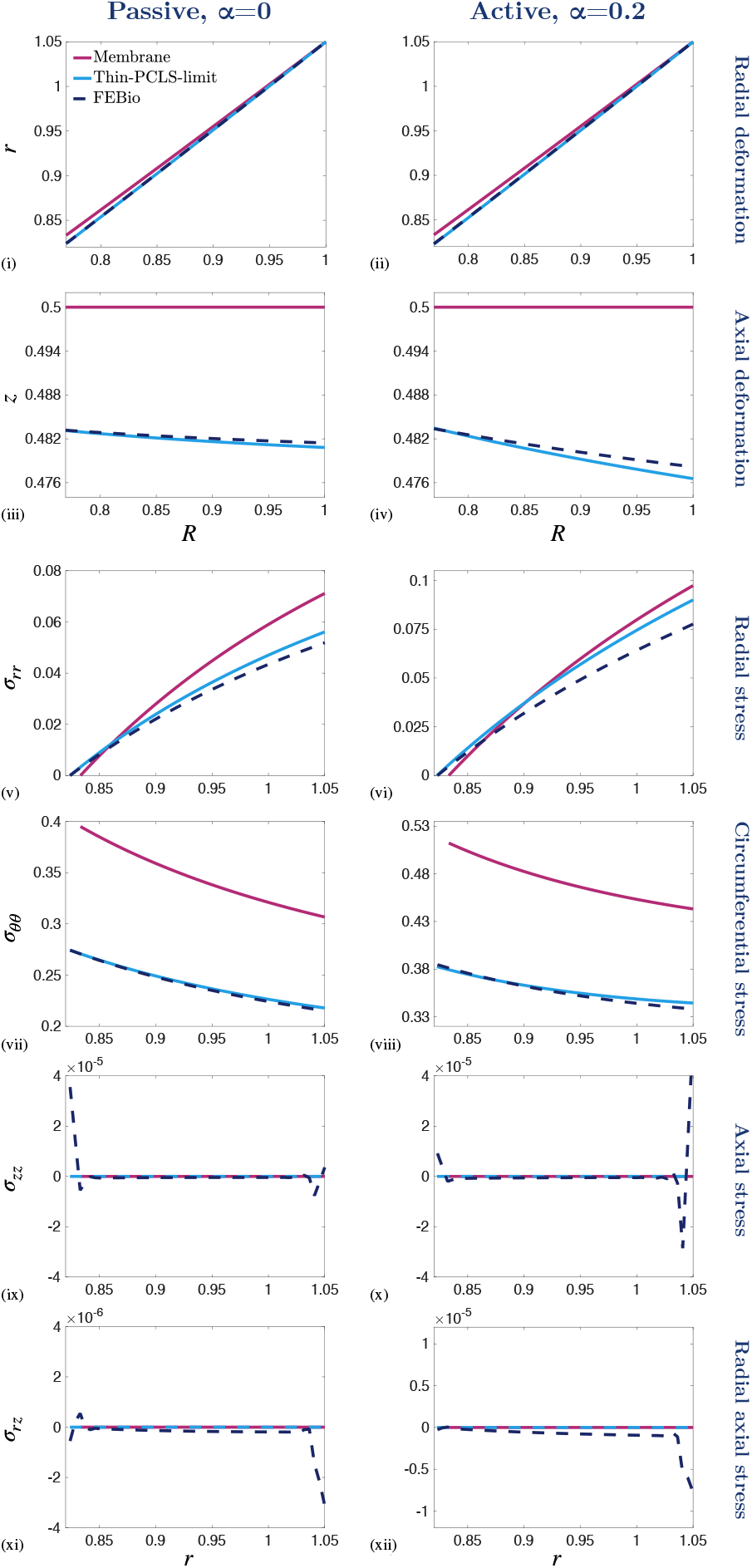
Comparison of the numerical simulations of the full model, with those of the membrane model, and the thin-PCLS-limit to demonstrate their validity. (i)–(ii) Radial displacement, *r*, (iii)–(iv) axial displacement, *z*, and (v)–(xii) Cauchy stress components, ***σ***, plotted as functions of undeformed radius, *R*, and deformed radius, *r*, at the undeformed centre of the PCLS (*Z* = 0), respectively. A 5% fixed stretch is applied to the PCLS in the passive, *α* = 0, (1^st^ column) and active, *α* = 0.2, (2^nd^ column) case. The aspect ratio *ε* = 0.01 throughout and the remaining parameter values are provided in Table 1 in Appendix C.

We find that, despite its relative mathematical and computational simplicity, the thin-PCLS-limit provides an excellent approximation to the full model, showing good quantitative agreement in all variables. In contrast, the membrane model is unable to replicate the full model behaviour. The one-dimensional geometry of the membrane approximation constrains the inner radius of the membrane to deform corresponding to the displaced outer boundary in order to preserve incompressibility. As a result, we observe an increased radial deformation and elevated radial and circumferential stress in the membrane approximation compared to the thin-PCLS-limit approximation and the full model (Figures 5 (i)–(ii), (v)–(viii), respectively). In contrast, the thickness of the PCLS in both the full model and the thin-PCLS-limit allows the generated stresses to be absorbed by the axial deformation (Figures 5 (iii), (iv)). Hence, the thin-PCLS-limit provides a more realistic representation of the full problem (for small *ε*) than the membrane.

Active contraction accentuates the radial deformation of the PCLS and the thickness of the PCLS decreases accordingly in order to maintain tissue incompressibility (*cf*. Figures 5 (iii), (iv)). This feature is only observed in the full model and the thin-PCLS-limit. Further contraction-induced deformation is not permitted in the membrane approximation due to the one-dimensional geometry and incompressibility constraint, and as a result, active contraction simply increases the stress generated in the membrane. Hence, there are significant qualitative and quantitative differences observed in the stress distributions of the two approximations and only the thin-PCLS-limit provides a suitable approximation to the full model.

The full model exhibits rapid variation in the axial and shear stress components near the inner and outer airway radii (Figures 5 (ix), (xii)). However, this (small-amplitude) boundary layer behaviour in the axial and shear stress components near the airway boundaries (that is evident in the full model) is not captured by either of the simple models. Although the amplitude of these effects is very small, we believe that these are not numerical artefacts as they span multiple elements. Therefore, a boundary layer analysis of these features forms a natural future extension of this work.

## 5 Thin-PCLS-limit parameter exploration

In the preceding numerical experiments, our parameter choices follow that of Hill et al. (2018) or are estimated from the literature. The parameter values are provided in Table 2 in Appendix C. In this section we take advantage of the computational tractability of our reduced model ((31), (34b) and (39)) to explore the influence of the airway’s mechanical properties on the model behaviour, and in particular, examine differences in the constituents’ stresses that cause TGF-*β* activation. Such parameter exploration is computationally prohibitive in the full model.

**Table 2.** Table of dimensionless parameter values used in the thin-PCLS-limit model in Section 5. Dashes denote the given baseline value. Where multiple values are given, the value used for each plot in the figure is specified in the figure’s legend. Where ranges of parameter values are given, the parameter is varied and takes all values in the range.

| Parameter | Baseline | Figures |  |  |
| --- | --- | --- | --- | --- |
|  |  | 6 | 7 | 8 |
| $R_{\text{in}}$ | 0.7692 | - | - | - |
| $R_{\text{out}}$ | 1 | - | - | - |
| $r_{\text{dis}}$ | $\{1, 1.05, 1.15\}$ | $1 \leq r_{\text{dis}} \leq 1.2$ | - | - |
| $\Phi_c$ | 0.2 | - | - | - |
| $\Phi_m$ | 0.1 | - | - | - |
| $\omega$ | 0.3492 | - | - | - |
| $\zeta$ | 1.48 | - | - | - |
| $\mu$ | $\{\frac{1}{3}, 1, 3\}$ | - | 3 | $0 \leq \mu \leq 4$ |
| $\alpha$ | 0 | - | $0 \leq \alpha \leq 4$ | 1 |

The effect of the imposed radial displacement on the constituent stress, in the absence of contraction, is illustrated in Figure 6. Here, we increase the fixed stretch applied from 0% (unstretched) to 20% (the maximum imposed in the experiments that are our primary motivation (Tatler, 2016)). Over this range, we observe a significant increase in stress heterogeneity, with high radial (circumferential) stresses evident at the outer (inner) airway wall (Figure 6). For both the ASM and ECM components, we see that the circumferential stress dominates over the radial stress (*e.g., cf*. Figures 6 (i), (iv) and (vii), (x)). This is to be expected in the passive case due to the strain-stiffening of the ECM that occurs in direction of the circumferential fibres when stretched. However, we see that increasing the stiffness of ECM relative to that of ASM (*μ*) dramatically alters the distribution of the constituent radial stress (*cf*. Figures 6 (i), (iii) and (vii), (ix)).

**Fig. 6.**
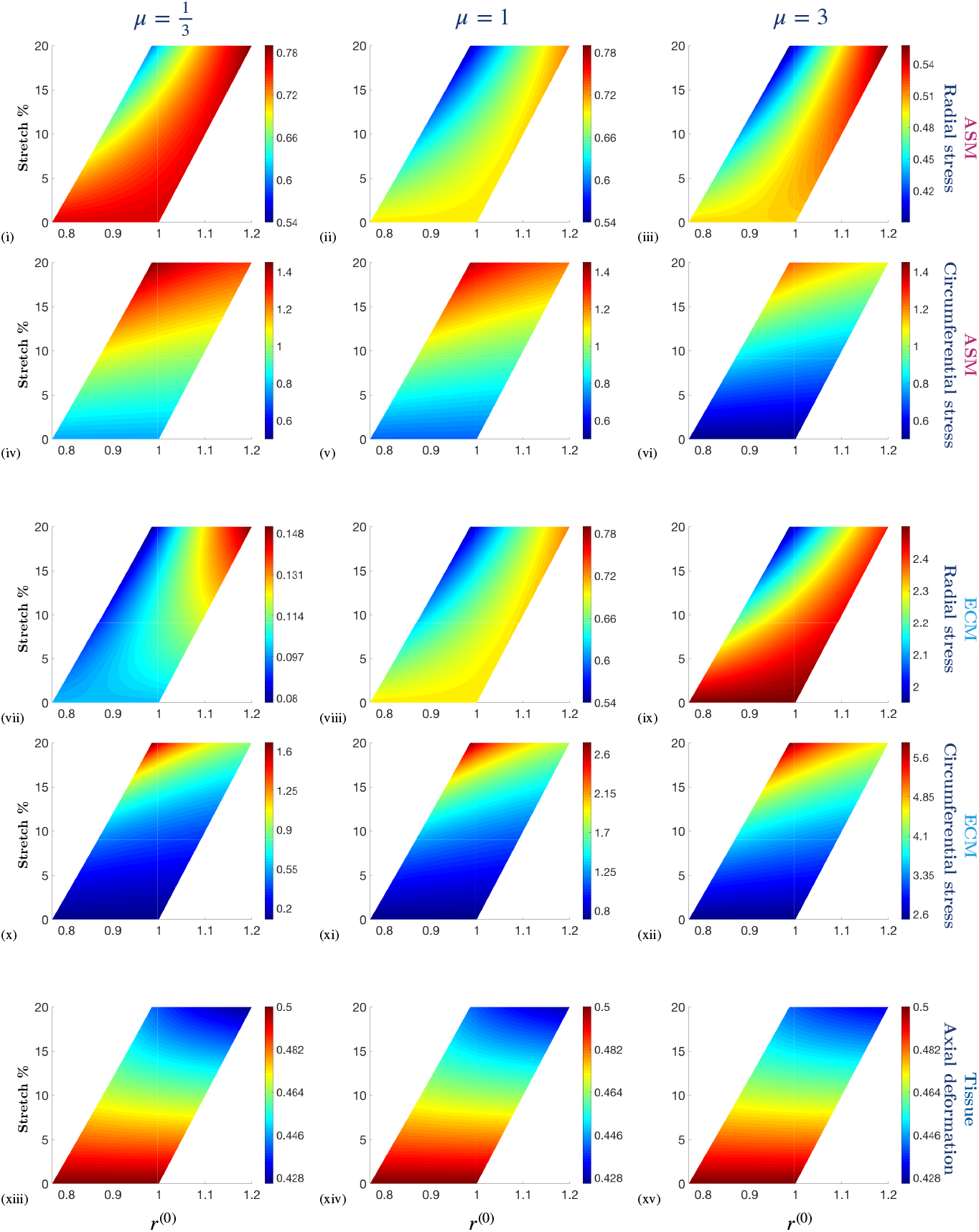
The effect of stretch (applied at the outer boundary of the PCLS) on the constituent Cauchy stress components, ***σ**_c_* and ***σ**_m_*, and the axial deformation at the upper surface, 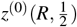, as a function of deformed radius, *r*^(0)^, for different stiffnesses of ECM relative to that of ASM, *μ*. Simulation parameter values are provided in Table 2 in Appendix C.

As the imposed radial stretch is increased from zero, the axial deformation becomes less uniform across the airway thickness (Figures 6 (xiii)–(xv)). Moreover, we observe that the heterogeneity of the axial thinning with increasing stretch is exaggerated with ECM compliance (*cf*. Figures 6 (xiii), (xv)). This suggests that the stiffness of the ECM provides resistance to the imposed stretch across the airway wall, in addition to the associated strain-stiffening. As a result, the profile of the PCLS is more uniform for stiff ECM and thicker (thinner) at the inner (outer) radius for compliant ECM. When the ECM is relatively compliant, the stress state of the ASM is higher than that of the ECM for small stretches (due to the passive isotropic material properties, *i.e. μ*) (*cf* Figures 6 (i), (vii)). However, strain-stiffening of the ECM increases exponentially with stretch and as a result, the stress state of the ECM increases more significantly with stretch than that of the ASM (*cf*. Figures 6 (iii), (ix)).

The influence of ASM contractility on the airway constituent stress and deformation, at fixed imposed stretch, is illustrated in Figure 7. As expected, increasing the contractile force generated by the ASM leads to significant radial contraction of the airway, and associated resistance to axial thinning at the inner radius (Figures 7 (xiii)–(xv)). Correspondingly, we observe elevated and increasingly non-uniform radial stress of the ASM and ECM constituents in Figures 7 (i)–(iii) and (vii)-(ix), respectively.

**Fig. 7.**
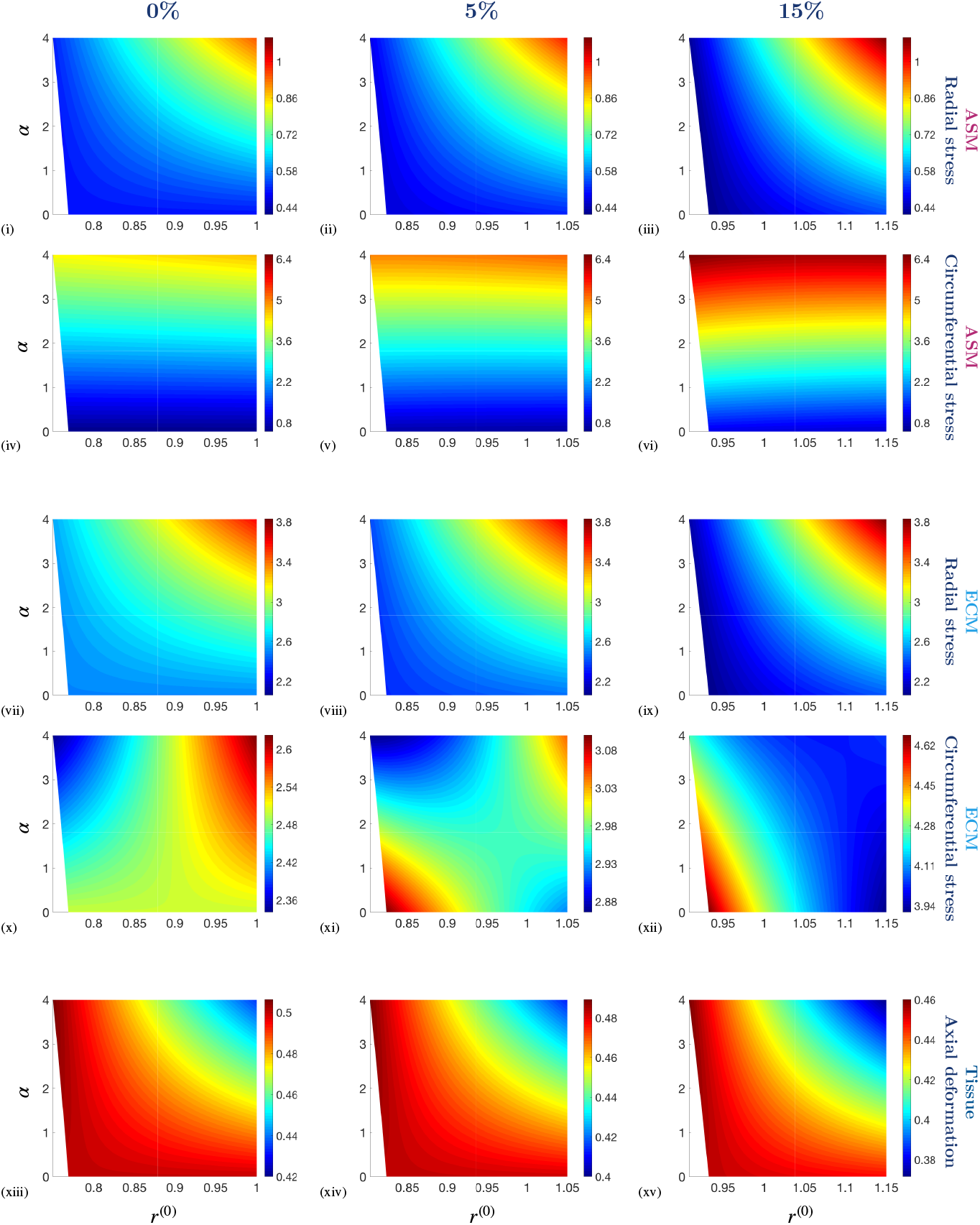
The effect of constant contractile force, *α*, on the Cauchy constituent stress components, ***σ**_c_* and ***σ**_m_*, and the axial deformation at the upper surface, 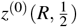, as a function of deformed radius, *r*^(0)^, in the presence and absence of fixed stretch applied at the outer boundary of the PCLS with amplitude 0%, 5% and 15%. Simulation parameter values are provided in Table 2 in Appendix C.

In general, the stress distributions are qualitatively similar at each amplitude of fixed stretch, with a small stress increase at a larger fixed stretch. The circumferential stress of the ECM is an exception to this general observation and displays significantly different heterogeneous distributions for each imposed fixed stretch (*cf*. Figures 7 (x), (xi) and (xii)). More specifically, in the absence of stretch, the circumferential stress of the ECM is maximal at the outer radius when the contractile force is high (Figure 7 (x)). In contrast, in the presence of a 15% stretch, the circumferential stress of the ECM is maximal at the inner radius and when there is no contractile force (Figure 7 (xii)). The transition between these two modes is evident in Figure 7 (xi).

In contrast to our previous observations for increasing stretch in Figure 6, we see that increasing the contractile force generated by the ASM leads to comparable radial and circumferential stress components of the ECM (Figure 7). Here, the increasing contractile force and the strain-stiffened ECM leads to the constituents being comparably stressed.

The influence of constituent stiffness on the airway wall stress and deformation, at fixed imposed stretch, is illustrated in Figure 8. The ratio of the passive isotropic stiffness of ECM to that of the ASM is given by *μ*. The ECM is more compliant than the ASM for *μ* < 1, stiffer than the ASM for *μ* > 1, and has the same stiffness as the ASM for *μ* = 1. Here, we increase *μ* in the presence of a constant contractile force generated by the ASM, *α*.

**Fig. 8.**
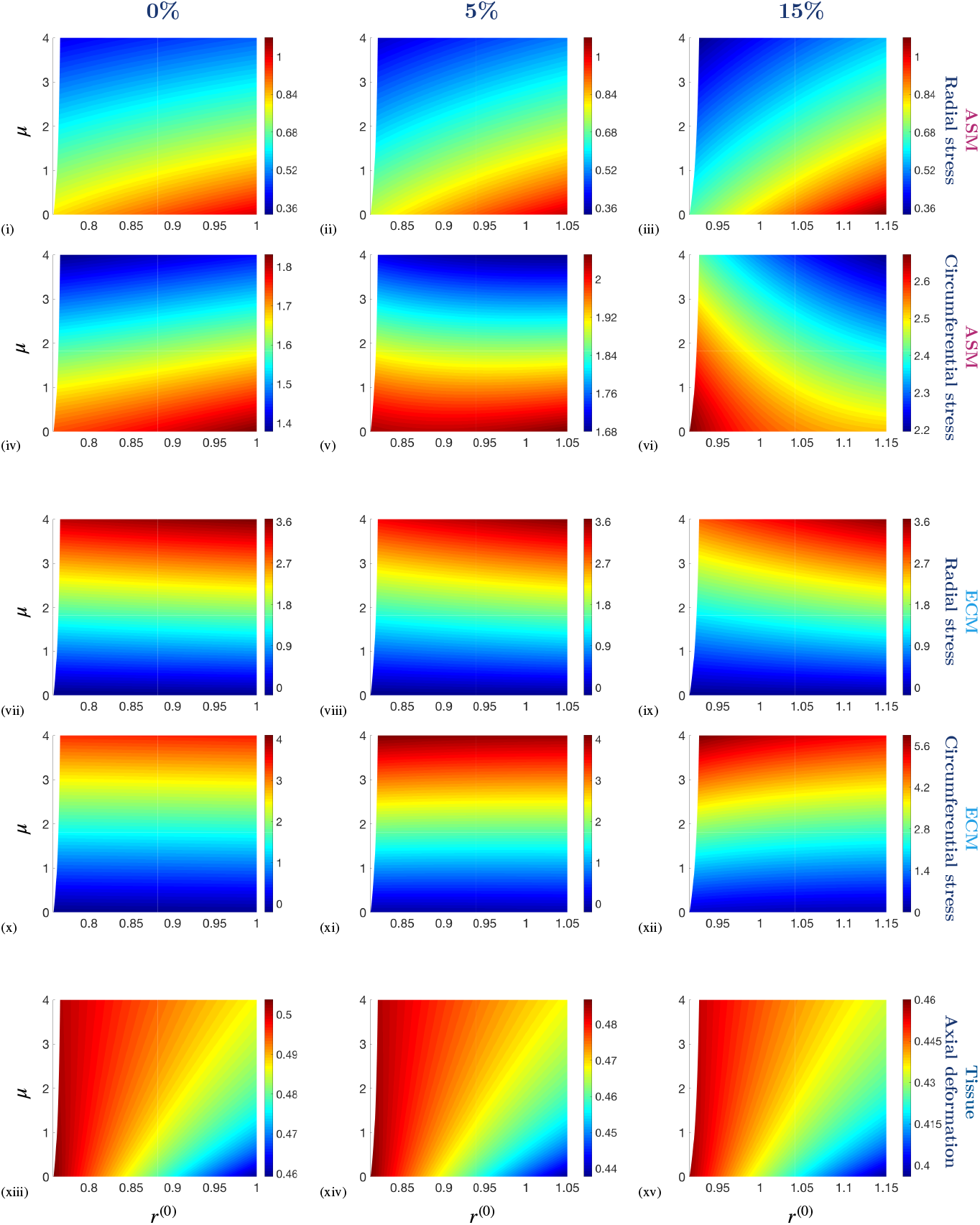
The effect of the stiffness of ECM relative to that of ASM, *μ*, on the constituent Cauchy stress components, ***σ**_c_* and ***σ**_m_*, and the axial deformation at the upper surface, 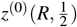, as a function of deformed radius, *r*^(0)^, in the presence and absence of a fixed stretch applied at the outer boundary of the PCLS with amplitude 0%, 5% and 15%. Simulation parameter values are provided in Table 2 in Appendix C.

When the ECM is more compliant than the ASM (*μ* < 1) we observe a slight reduction in radial contraction compared to the case for which the ECM stiffness exceeds that of the ASM (*μ* > 1). Correspondingly, there is an increased resistance to axial thinning with increasing *μ* observed in Figures 8 (xiii)–(xv). As a result, we see that the magnitude of the stress components of the ASM decrease with increas-ing *μ* (due to the decreased radial contraction), whilst the magnitude of the stress components of the ECM increase with increasing *μ* (*cf*. Figures 8 (v), (xi)). These observations persist and are emphasised under the application of fixed stretch (due to additional strain-stiffening of the ECM when stretched).

In general, we see that the stress distributions of the ASM display greater nonuniformity across the airway radius than that of the ECM, particularly when stretched. For example, the circumferential stress of the ASM and ECM differ significantly (*cf*. Figures 8 (vi), (xii)). As the stiffness of the ECM increases, the circumferential stress of the ASM remains higher at the outer radius than at the inner radius in the absence of stretch (Figure 8 (iv)). However, in the presence of a 15% stretch, the circumferential stress of the ASM varies significantly across the airway wall and is higher at the inner radius and lower at the outer radius (Figure 8 (vi)). Therefore, imposed stretch induces a dramatic change in the distribution of the circumferential stress of the ASM and the apparent transition between these two extremes is observed in Figure 8 (v). This behaviour is similar to that exhibited by the ECM when increasing the contractile force generated by the ASM in Figures 7 (x)–(xiii).

## 6 Discussion

Despite its prevalence in the population, the causes of asthma remain poorly understood; in particular, the feedback mechanisms linking inflammation, bronchoconstriction and cytokine activation are yet to be elucidated. To help address this, we develop a nonlinear fibre-reinforced biomechanical model of an airway in PCLS, an *ex vivo* assay widely used for studying asthmatic airway biomechanics. Our model accommodates agonist-induced ASM contractility and ECM strain-stiffening and allows us to examine the stress distributions of these individual constituents within the airway wall.

Direct numerical simulation of the model by means of the FEBio software (Maas et al., 2012) reveals the internal stress state of an axisymmetric airway within a PCLS under imposed deformation, and highlights the distinct qualitative and quantitative differences induced by ASM contraction. Such information is of key importance in interpreting PCLS experiments, and in particular those that seek to understand the above described feedback mechanisms. However, the computational complexity of this model precludes thorough investigation of the parameter space, mathematical analysis or coupling to time-dependence. To address this, we consider two reductions of the full model. First, we adopt a membrane representation, where axial variation is neglected *a priori*, and the PCLS thickness remains unchanged upon deformation. Secondly, and in view of the typical dimensions of the PCLS and numerical evidence, we consider an asymptotic reduction, appropriate for the limit in which the PCLS thickness is much smaller than the typical airway radius, and in which we are able to retain a description of the radial and axial airway deformation and the associated stresses. In each case, we reduce the model to one spatial dimension; the membrane model admits analytical solutions, while in the thin-PCLS-limit the model reduces to a pair of coupled nonlinear ODEs describing the deformation, numerical solutions to which are obtained via a shooting method. We find that the membrane model is unable to capture the full model behaviour, but that the asymptotically-reduced model provides an excellent approximation to the full model, at reduced computational cost.

Crucially, this computationally tractable model that we have developed allows for comprehensive investigation of the mechanisms underpinning pro-remodelling and contractile cytokine activation in asthmatic airways, a key aspect of the pathogenesis and presentation of asthma, that has only recently received attention. In particular, our future work will consider the positive mechanotransductive feedback loop between airway contraction and activation of the TGF-*β*, that is implicated in long-term remodelling. Other important future considerations include developing our asymptotic reduction to accommodate, for example, spatially-dependent airway composition data (Brook et al., 2019) and undertaking parameterisation and model validation. In addition, as the PCLS thickness is reduced, we observe rapid variations in the axial and shear stress components near the inner and outer airway radii that are not captured by the asymptotic (or membrane) model; a boundary layer analysis of these features forms a natural extension of this work. More advanced theoretical considerations include development of a model accommodating unconstrained multiphase solid mechanics. This will allow for the differing strain rates of the two constituents to be examined, which may be important in understanding the cell-mediated activation of cyotkines in the PCLS.

## A Numerics

Numerical solutions to the full biomechanical model (equations (20) and (21)) are obtained via a finite element method, implemented in the software FEBio (Maas et al., 2012). To confirm accuracy of our results, we carry out suitable mesh convergence tests by uniformly refining the mesh. An example of the meshed geometry is illustrated in Figure 9. We demonstrate that our results converge to a solution with increasing number of elements in Figure 10.

**Fig. 9.**
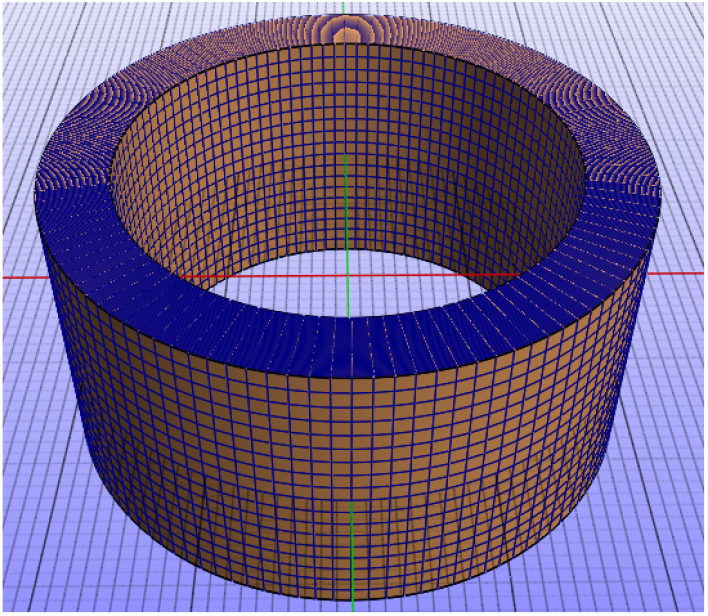
Example of the meshed geometry of an airway in the PCLS with aspect ratio *ε* = 1. Mesh produced in the software suite PreView (complementary to FEBio (Maas et al., 2012)).

**Fig. 10.**
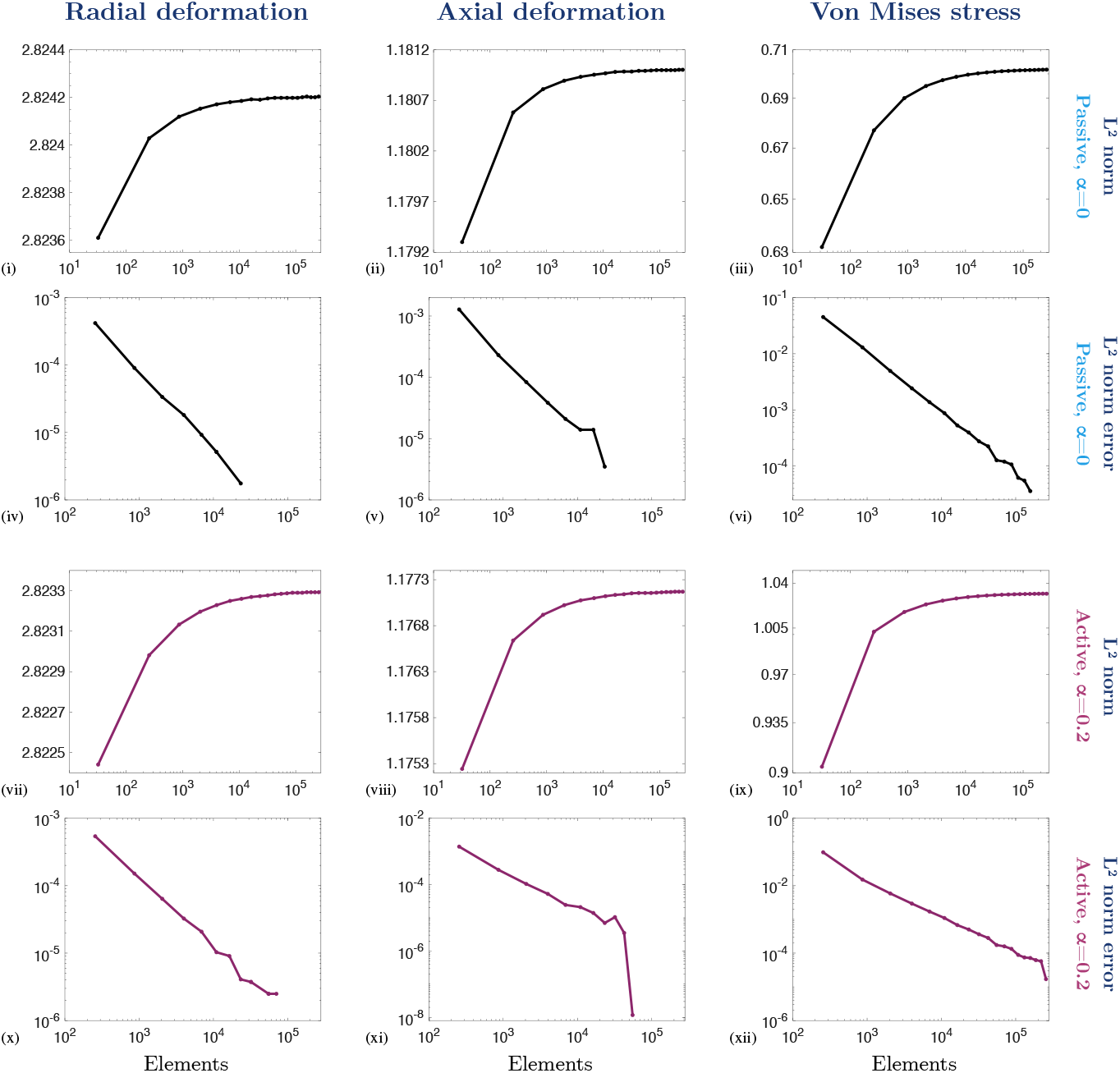
Mesh convergence tests. (i)–(iii) The *L*^2^ norm (40) and (iv)–(vi) the *L*^2^ norm error between iterations of mesh refinement (42) of the radial deformation, *r*, axial deformation, *z*, and Von Mises stress, *σ*_VM_, in the passive, *α* = 0, case, plotted against the number of elements in the mesh. (vii)–(ix) The *L*^2^ norm (40) and (x)–(xii) the *L*^2^ norm error (42) of *r, z* and *σ*_VM_ in the active, *α* = 0.2, case, plotted against the number of elements in the mesh. A fixed 5% stretch is applied with *ε* = 1 throughout and the remaining parameter values are provided in Table 1 in Appendix C.

The *L*^2^ norm of a *q* × *s* matrix **A** is defined as

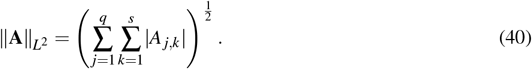

We use the *L*^2^ norm (40) as a metric in our mesh convergence tests in order to provide a global representation of the change in solution. In addition to the radial and axial deformation variables, *r* and *z*, we consider the Von Mises stress, *σ*_VM_, given by

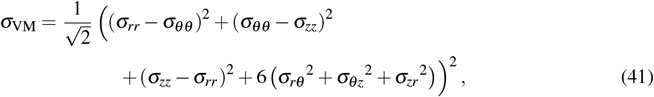

in order to provide an overall representation of the Cauchy Stress components. For consistency, ||*r*(*R, Z*)||_*L*_^2^, ||*z*(*R, Z*)||_*L*_^2^ and ||*σ*_VM_(*R, Z*)||_*L*_^2^ are evaluated at the nodal positions of the coarse mesh at each iteration and are plotted against the number of elements in the mesh to show that the solutions converge appropriately both in the passive and active contraction case (Figures 10 (i)–(iii) and (vii)–(ix), respectively).

Using (40), we define the *L*^2^ norm error between iterations, *n*, as

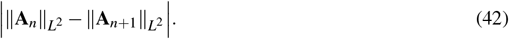

We show that the errors between iterations decay to zero in Figures 10 (iv)–(vi) and (x)–(xii), again in both the passive and active contraction case respectively. Note that we have carried out mesh convergence tests for *ε* ∈ {0.01,0.1,1}, with and without a prescribed stretch, but limit the results displayed in Figure 10 to *ε* = 1 with a 5% fixed stretch applied for concision.

## B Thin-PCLS-limit model reduction

Guided by the numerical evidence (provided in Section 3 and below), in this section we demonstrate that the general expansions, for which *r* = *r*(*R, Z*) and the leading order term for 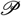 is 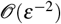 (to obtain a proper leading order balance in the Cauchy stress) can be reduced to that shown in (29).

The general asymptotic expansions for each of the deformation variables are as follows,

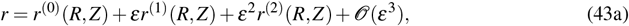

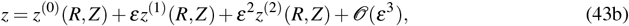

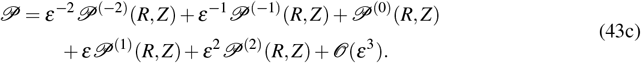

As outlined in Section 4.2, we assume that *Φ_c_, Φ_m_, μ, ω, ζ* and *α* are all 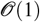 constants in the strainenergy function for the whole tissue, *W* (18c). Therefore, the following derivatives of the strain-energy functions are 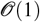 constants: *W*_*c*1_, *W*_*m*1_ and *W*_1_. The derivative of the strain-energy function with respect to the fourth invariant, *W*_4_, is a function of radius, *r*. Therefore, using the asymptotic expansions (43) and expanding for small *ε* in *W*_*m*4_, we obtain the leading order terms in the derivative of the strain energy-function, with respect to the fourth invariant, for each constituent

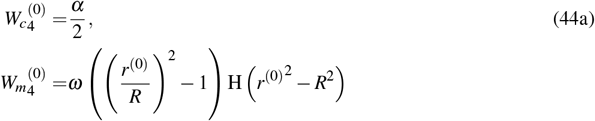

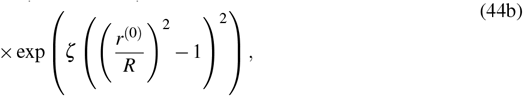

and correspondingly for the whole tissue

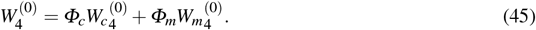

At the next order, 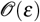, we obtain

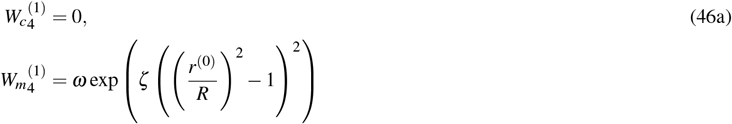

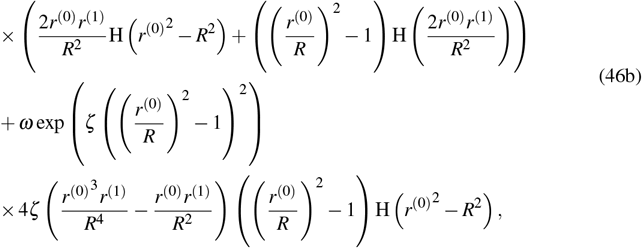

and correspondingly for the whole tissue

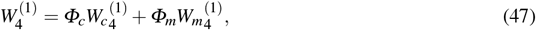

where, as previously, the notation 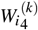 for *i* ∈ {*c,m*} refers to the 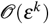 term in *W*_*c*4_ and *W*_*m*4_. Using the expansions (43) and expanding for small *ε* in (20b)-(20c)), at the leading order, 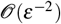, the governing equations (20b)–(20c) read

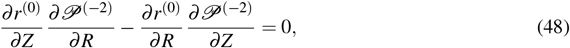

subject to the following free boundary conditions at the upper, lower and inner surfaces of the PCLS (21b):

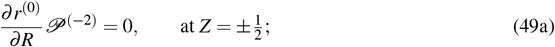

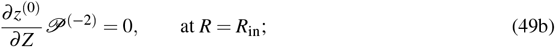

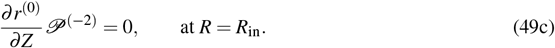

At the next order, 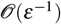, the governing equations (20b)–(20c) read

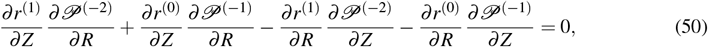

subject to the free boundary conditions (21b)–(21d):

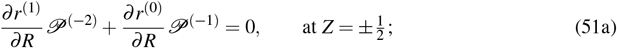

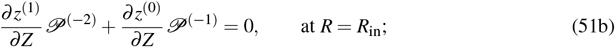

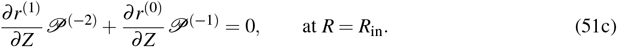

A solution to (48) that satisfies the free boundary conditions (49) is 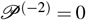, and thus, a solution to (50) that satisfies the boundary conditions (51) is 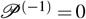. This solutionis consistent with our numerical solution obtained in FEBio in Section 3 (*e.g*. Figure 2), where the pressure is an 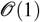 quantity, *i.e*. where 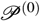 is the leading order term. Subsequently, at 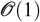, the governing equations (20) read

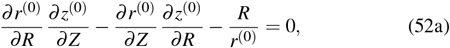

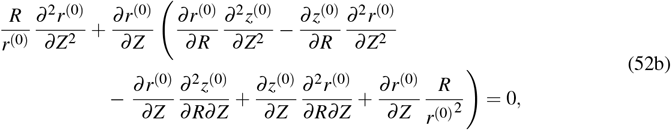

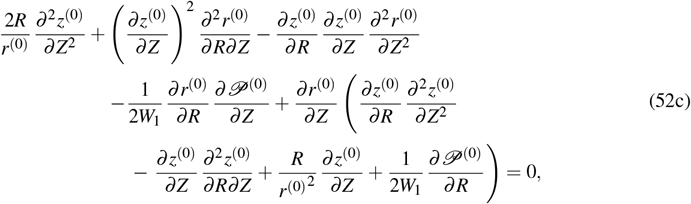

subject to the displacement boundary condition at the outer radius (21a),

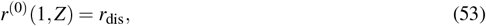

and the free boundary conditions (21b)–(21d):

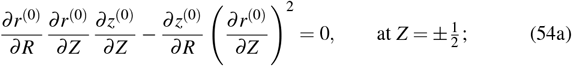

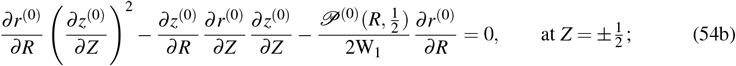

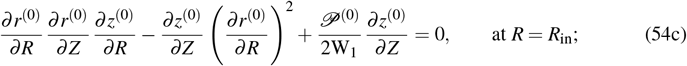

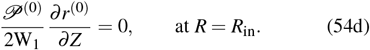

We seek to obtain (30) which requires *r*(0)(*R*). We note that *r*^(0)^(*R*) is a solution to (52) that satisfies the displacement boundary condition (53) and the free boundary conditions (54). Moreover, numerical evidence provided in Figures 11, 1^st^ column (illustrating the effect of reducing the aspect ration, *ε*, on the first derivative of the radial deformation, *r*, with respect to *Z*) shows that the dependence of *r* on *Z* is very weak. In view of this, we relegate *Z* dependence to the next order, and the governing equations (52) reduce to

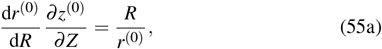

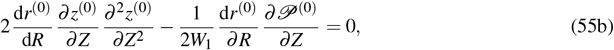

and the free boundary conditions (54) reduce to:

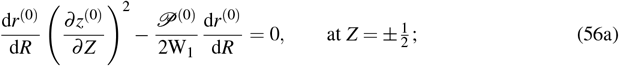

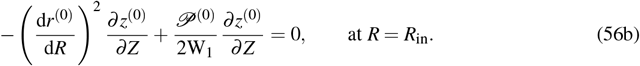

as previously in Section 4.2.

**Fig. 11.**
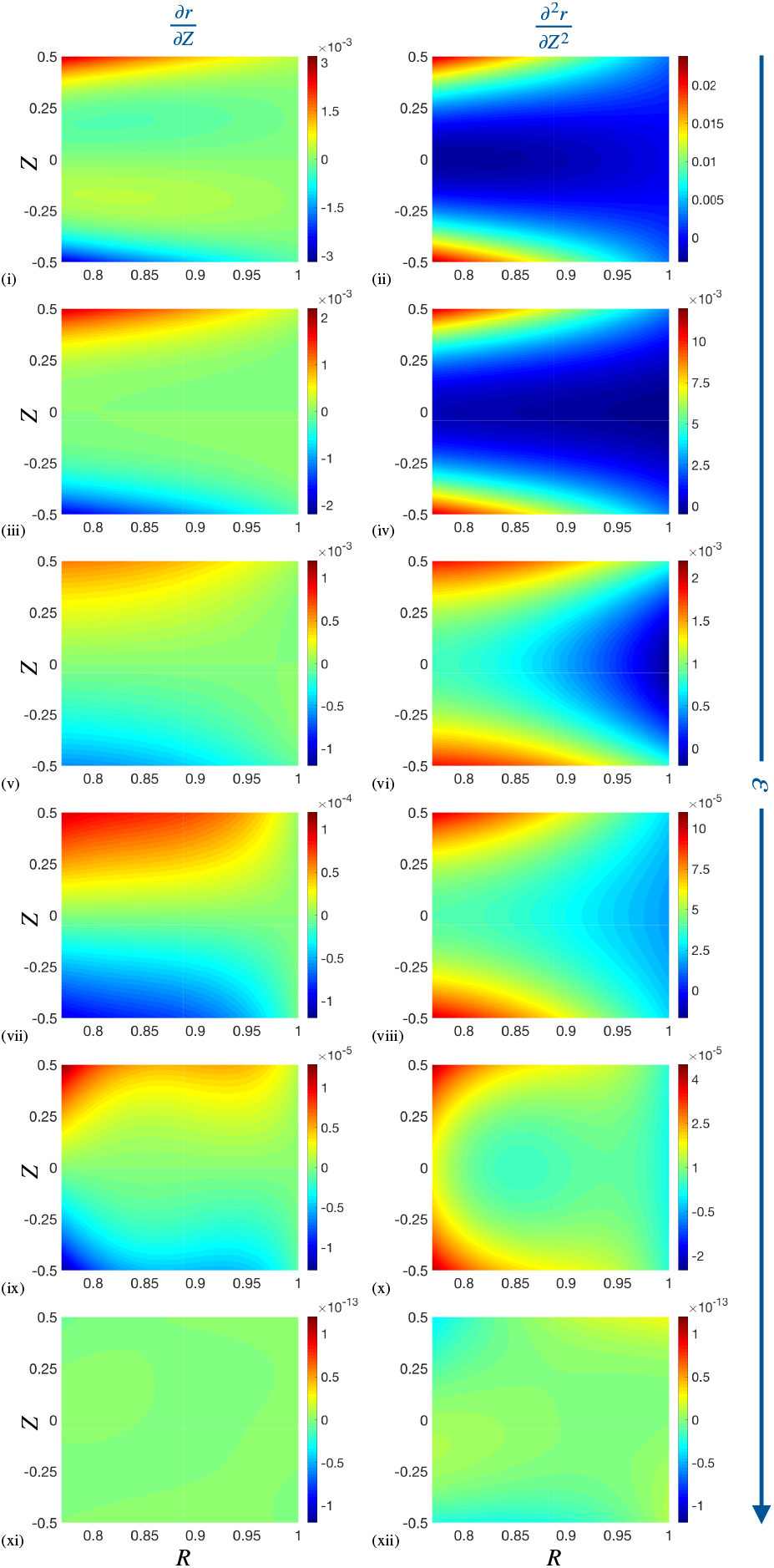
The effect of reducing the aspect ratio, *ε*, on the first derivative (1^st^ column) and second derivative (2^nd^ column) of the radial deformation *r* with respect to *Z*, in the passive case, *α* = 0, and under the application of a 5% fixed stretch. Results plotted in the undeformed reference configuration (*R, Z*). *ε* decreases in the direction of the arrow for *ε* ∈ {1,0.5,0.25,0.1,0.025,0.01}. The remaining parameter values are provided in Table 1 in Appendix C.

From (55a), we note that

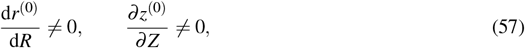

and rearrange the reduced boundary conditions (56) to obtain

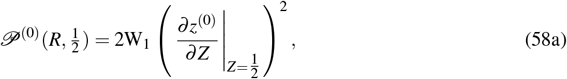

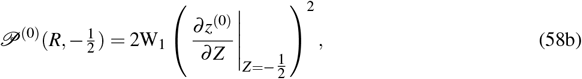

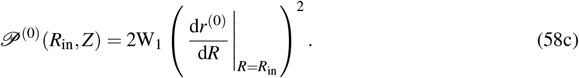

The boundary condition (58c) holds for all *Z*; however, *r*^(0)^ is independent of *Z*, and therefore 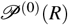. Subsequently, the 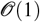 governing equations (55) reduce to

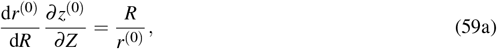

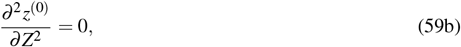

and provide

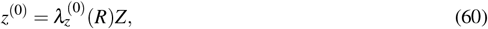

where the arbitrary function of *R* arising from the integration of (67) vanishes due to axial symmetry in *z*^(0)^ about the axial centre line, *Z* = 0, as previously in (33). Substituting (60) into (58) and (59) we obtain

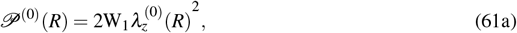

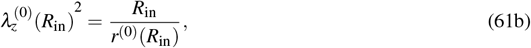

as previously in (34).

So far we have demonstrated that *r*^(0)^(*R*) and 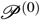 are appropriate leading order approximations in (29); we now show that *r*^(1)^(*R*) and 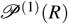 follow.

Using the preceding information, and in particular (57), at 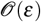 the governing equations read

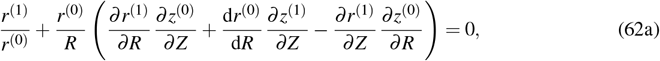

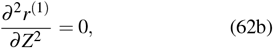

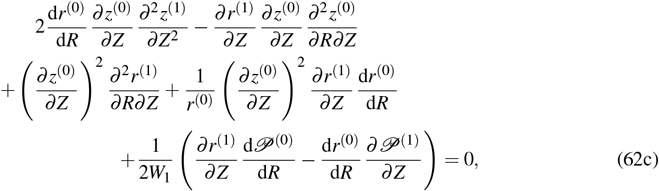

subject to the displacement boundary condition at the outer radius (21a),

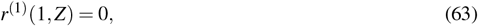

and the free boundary conditions (21b)–(21d):

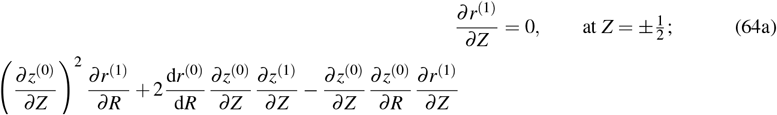

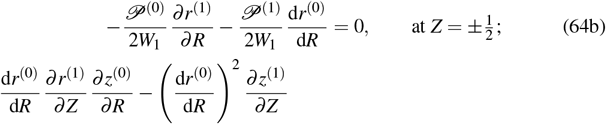

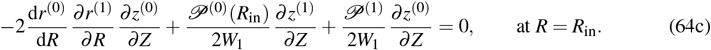

The 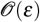 governing equation (62b) together with the boundary condition (64a) provides *r*^(1)^(*R*), and hence, the 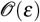 governing equations (62) reduce to

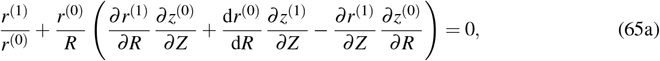

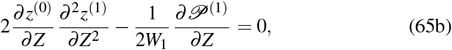

and, using (61), reduces the 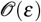 free boundary conditions (64) to

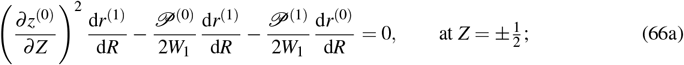

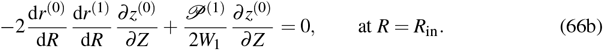

The free boundary condition (66b) holds for all *Z*; however, all terms except 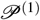 are independent of *Z*. Therefore, we conclude, 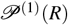. Subsequently, the 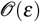 governing equations (65) reduce to

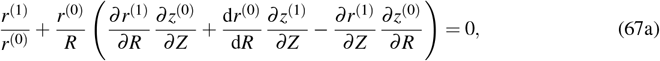

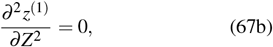

and provide

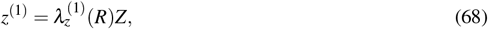

where, again, the arbitrary function of *R* arising from the integration of (67b) vanishes due to the symmetry in *z*^(0)^ about the axial centre line, *Z* = 0. Substituting (60) and (68) into the free boundary conditions (66) we obtain

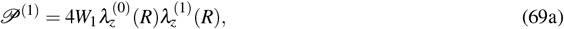

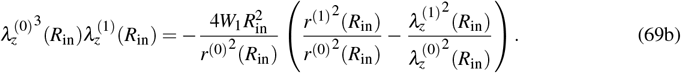

To summarise, we have reduced the problem to two leading order variables; the radial deformation, *r*^(0)^(*R*), and the axial stretch, 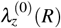. The governing equations together with the free surface boundary conditions up to 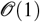 provide the pressure 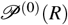 in terms of *r*^(0)^(*R*) and 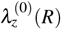. The leading order governing equations enforce incompressibility (59a) subject to the displacement boundary condition at the outer radius (53) and the free surface boundary conditions via a relation at the inner radius in (61b). The governing equations together with the free surface boundary conditions up to 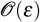 provide 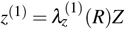 in (68) and 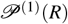 in (69a); however, we require a leading order equation that governs the balance of linear momentum to determine the two variables, *r*^(0)^(*R*) and 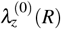. Therefore, at 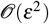 the linear momentum equations read

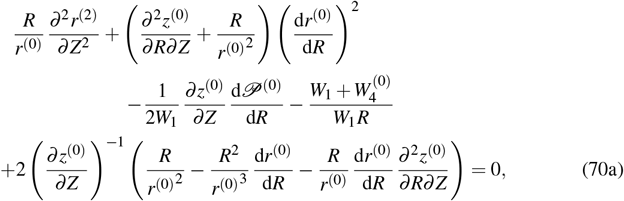

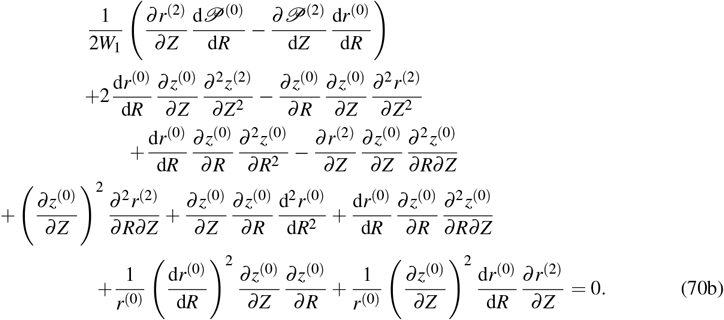

Using (70a), we seek to obtain (38) in order to close the leading order problem. As previously, equation (70b) introduces higher order terms (i.e. *z*^(2)^ and 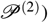 which are not of interest for the leading order problem and is therefore not needed here.

Numerical evidence provided in Figures 11, 2^nd^ column shows that the dependence of *r* on *Z* is weak and importantly, that 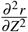 is much smaller than the other leading order quantities in (70a). As a result, the remaining 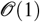 terms in (70a) dominate to reveal (38). Moreover, for *ε* = 0.01, Figures 11 (xi) and (xii) suggest that *r*^(2)^ = *r*^(2)^(*R*) and support our expansions in (29). Subsequently, substituting (60) and (61a) into (70a) provides an equation for 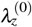 and hence, together with (59a), we obtain the pair of coupled ODEs (39). As outlined in Section 4.2, the coupled ODEs (39) together with the boundary condition (53) and the relation (69b) provides a boundary value problem that describes the leading order radial and axial deformation, and from which the leading order Cauchy stress components for the whole tissue and each of the constituents follow directly. We note that the boundary condition (69b) on 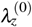 is posed at the (unknown) deformed inner radius. We therefore solve (39) numerically by treating 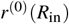 as a shooting parameter and seek the solution set 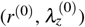 at *R*_in_ that satisfies (39), (53) and (69b).

In summary, we have demonstrated that the full biomechanical model of the PCLS may be suitably reduced in the thin-PCLS-limit (Section 4.2) via the asymptotic expansions (29) to obtain a leading order boundary value problem.

## C Parameter values

